# Resetting the yeast epigenome with human nucleosomes

**DOI:** 10.1101/147264

**Authors:** David M. Truong, Jef D. Boeke

## Abstract

Humans and yeast are separated by a billion years of evolution, yet their conserved core histones retain central roles in gene regulation. Here, we “reset” yeast to use core human nucleosomes in lieu of their own, an exceedingly rare event which initially took twenty days. The cells adapt, however, by acquiring suppressor mutations in cell-division genes, or by acquiring certain aneuploidy states. Robust growth was also restored by converting five histone residues back to their yeast counterparts. We reveal that humanized nucleosomes in yeast are positioned according to endogenous yeast DNA sequence and chromatin-remodeling network, as judged by a yeast-like nucleosome repeat length. However, human nucleosomes have higher DNA occupancy and reduce RNA content. Adaptation to new biological conditions presented a special challenge for these cells due to slower chromatin remodeling. This humanized yeast poses many fundamental new questions about the nature of chromatin and how it is linked to many cell processes, and provides a platform to study histone variants via yeast epigenome reprogramming.

**Highlights:**

- Only 1 in 10^7^ yeast survive with fully human nucleosomes, but they rapidly evolve
- Nucleosome positioning and nucleosome repeat length is not influenced by histone type
- Human nucleosomes remodel slowly and delay yeast environmental adaptation
- Human core nucleosomes are more repressive and globally reduce transcription in yeast

## Introduction

Because they serve as a central interface for hundreds of other proteins, histones are among the most conserved genes in eukaryotes (Talbert and Henikoff, 2010). They serve central cellular roles by regulating genome access, DNA compaction, transcription, replication, and repair (Talbert and Henikoff, 2017). Whereas higher eukaryotes evolved myriad histone variants with specialized functions, *Saccharomyces cerevisiae* (budding yeast) encodes but a few, a simplicity that has facilitated many fundamental discoveries in chromatin biology (Rando and Winston, 2012). But this begs the question: why do budding yeast have such streamlined chromatin compared to humans, and do differences in histone sequences reflect functional divergence (Figure 1A)? Might the simple yeast serve as a “chassis” for understanding how histone variants exert control over cellular transcription?

**Figure 1.**
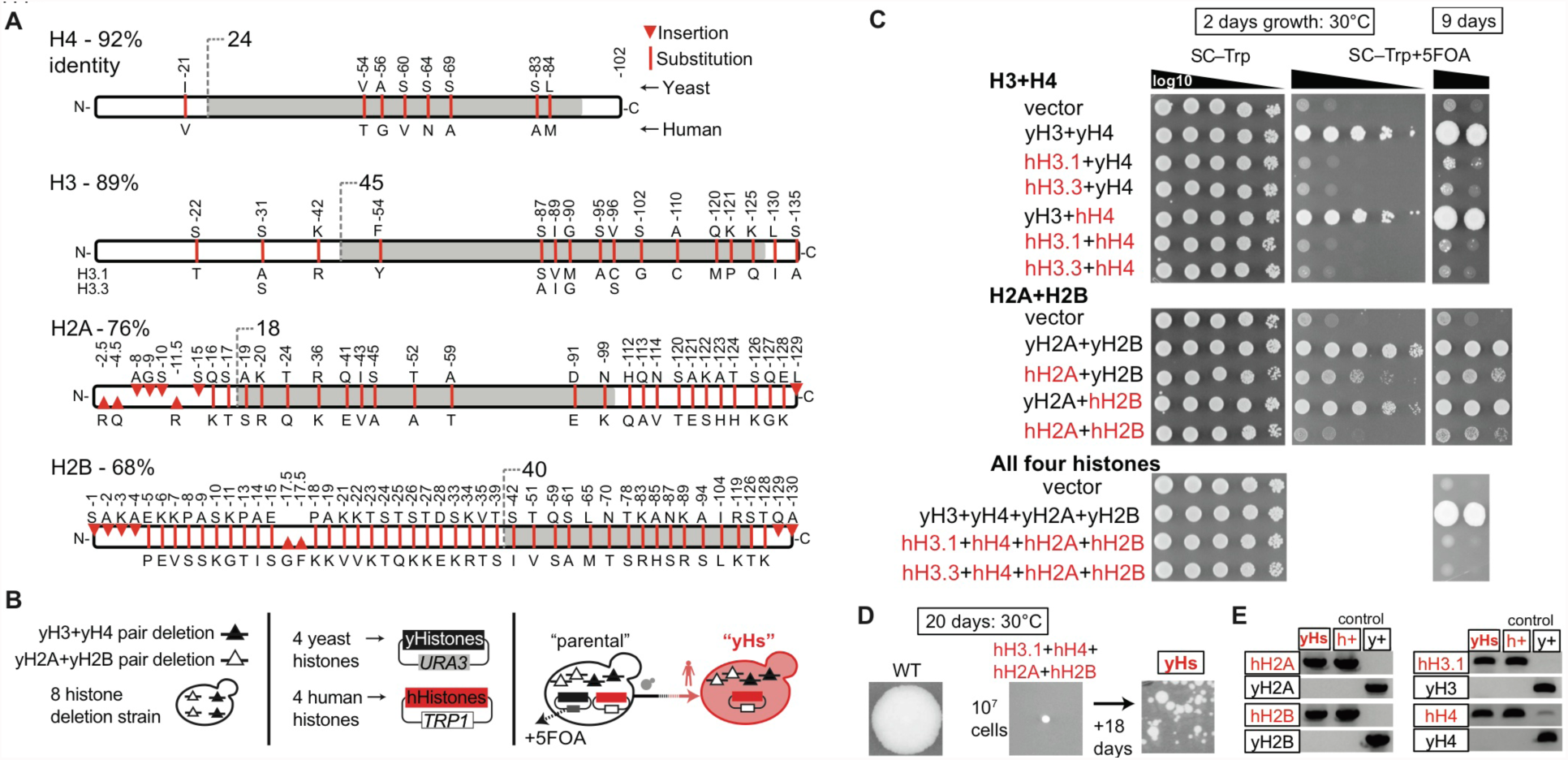
*Saccharomyces cerevisiae* can subsist on completely human core nucleosomes. (A) Human and budding yeast histones share from 68 to 92% protein identity. Red bars indicate residue positions that differ between the two species. Numbers refer to the yeast histones. Gray colored regions show the globular histone domains, and the white regions show the N- and C-terminal tails. (B) Dual-histone plasmid shuffle strategy (also see Figure S1). (C) The “humanization frequency” by which each human histone gene in the context of the relevant histone pairs or all four histone genes complement deletions of the respective yeast counterpart was assessed as in (B). Spots show yeast 10-fold serial dilutions. The densest spot contains 10^5^ yeast. (D) Yeasts with completely human nucleosomes arise after 20 or more days on plates. Only 8 colonies have been isolated thus far, referred to as “yHs”. (F) PCRtag analysis of humanized yeast (Mitchell et al., 2015), and confirmed by sequencing extracted plasmids. The human histones differ in DNA sequences enough to enable straightforward PCR genotyping.

Here we asked a simple but far reaching question: to what extent can *S. cerevisiae* utilize the core histones from humans? Thus far, only single yeast genes have been individually humanized (Hamza et al., 2015; Kachroo et al., 2015; Laurent et al., 2016; Osborn and Miller, 2007), but never a whole protein complex, nor one of such central importance.

These studies could reveal intrinsically different properties between yeast nucleosomes and those from humans. We initially suspected two possible outcomes for our study: *i*) that human histones would fully complement in place of yeast histones because of their high conservation, suggesting that histone sequence divergence provides only minor functional differences; or *ii*) that human histones in yeast would very poorly complement or even fail entirely, suggesting that the divergent residues are highly optimized for each species and serve specialized or novel functions. Our results are consistent with the latter.

The humanized yeast had pronounced delays in adapting to new environmental conditions, likely due to slowed remodeling of human core nucleosomes. In addition, our results suggest that human core nucleosomes may have evolved to occupy DNA more tenaciously, as we observed reduced RNA content and greater DNA occupancy by MNase-seq. Finally, our results suggest that yeast may maintain human chromatin even when given access to native yeast histones. This may represent a type of chromatin “memory”, whereby cells partition and reproduce parental chromatin to new daughter cells. Thus, while the species-specific coevolution of histones and their associated protein networks is extensive, it is nonetheless possible to reprogram the epigenome of at least one organism to accept histones of a very distant relative.

## Results

### *Saccharomyces cerevisiae* can subsist on fully human core histones

*S. cerevisiae* cells possess a relatively simple repertoire of histones. The core nucleosome, the four histones H3, H4, H2A, and H2B, comprise duplicate copies in the genome – each with divergent promoters and terminators – for a total of eight histone copies (Eriksson et al., 2012) (Figure S1). Additionally, there are three histone variants, H2A.Z, CENPA, and H1 (*HTZ1, CSE4,* and *HHO1*, respectively), which were not altered here.

To humanize the core nucleosome of yeast, we constructed three strains in which the target histones of interest (e.g., all 8 core histones) were deleted from the genome. We then used a plasmid shuffle approach (i.e., yeast vs. human histones on plasmids) for quickly eliminating the native yeast histones in favor of their human counterparts, by 5-FOA counter-selection (Boeke et al., 1987) (Figures 1B and S1; *see Methods*). The dual plasmids, containing either human or yeast histone genes, are expressed from different sets of endogenous histone promoters and terminators to eliminate recombination. Any designed humanized histone strain will carry only half the target histone copies (e.g., 4 instead of 8 histones).

First, we determined the relative “humanization frequencies” (i.e., humanized colonies per cell plated) for individual or pairs of human histones. Human histone H4 (hH4) had the highest humanization frequency (fast growth, 20%), followed by hH2B (slow growth, 20%), hH2A (slow growth, 10^-2^), and finally hH3.1 and hH3.3 (very slow growth, 10^-4^ and 10^-5^ respectively) (Figure 1C). Combining hH4 with either hH3.1 or hH3.3 also produced very slow growth (frequency of 10^-4^ and 10^-5^, respectively), whereas combining hH2A with hH2B led to slow growth, but a modest humanization frequency of ~10^-3^. For both human histone combinations, we confirmed the loss of yeast histones by PCRtag analysis (Mitchell et al., 2015), which uses PCR to discriminate between sequence differences of yeast and human histones, and a lack-of histone mutations by Sanger sequencing of recovered plasmids (Figure S1D, E).

We then attempted to exchange all four histone genes simultaneously (Figure 1C). An “isogenic-WT” strain (yDT67) that replaces one native yeast histone plasmid with a plasmid containing the other set of native histones “shuffled” readily (Figures 1C and S1). Yet neither plasmid encoding fully human core nucleosomes containing either hH3.1 or hH3.3 produced colonies within nine days. However, upon plating at least 10^7^ cells and waiting 20 days we did see colonies representing humanized yeast, but only for the hH3.1 plasmid (Figure 1D). These humanized yeast colonies were confirmed via PCRtag analysis and sequencing of extracted plasmids (Figure 1E), which showed no mutations in the human histones. These humanized colonies are unlikely to contain residual yeast histones, as “old”-histones turn-over by at least two-fold per cell division (Annunziato, 2005; Radman-Livaja et al., 2011). Since each haploid cell contains about 67,000 nucleosomes (Brogaard et al., 2012), and a small yeast colony contains at least 10^7^ cells, those underwent at least 23 cell divisions. Therefore, the cells contain on average 0.01 original yeast nucleosomes, assuming an infinite nucleosome/histone half-life.

To date, we have only identified 8 such confirmed direct humanization events (“yHs”-series; Figure S1F). Increased (high-copy plasmid) or decreased (genomic integration) human nucleosome gene copy number did not enhance humanization frequency (Table S3). The “yHs” cells grow on both synthetic complete (SC) and yeast complete (YPD) medium at 30°C and 25°C, but not at higher temperatures (e.g., 37°C), can mate, and grow to various degrees on media that enhance defects in DNA replication, DNA repair, and vacuole formation (Figure S2A, B). Finally, each of the humanized strains possessed differing rates of substantially slower than normal growth, and frequently produced larger and faster growing colonies over time (Figure 1D and Figure S1H). These observations are consistent with the accumulation of suppressors. A second factor reducing the humanization frequency is that a substantial proportion of the humanized cells in a population are unable to form a living colony.

### Bypass of cell-division genes promotes growth with human nucleosomes

We performed evolution experiments on seven “yHs” lineages to determine how and to what extent yeast cells adapt to “live with” core human nucleosomes. The seven lineages were selected by serially diluting and sub-culturing liquid stationary phase cultures for 5 cycles (Figure 2A). The evolved pools and isolates outperformed the pre-evolved strains on solid media, and doubled 33% more rapidly in liquid culture (Figures 2B and S2C). We then performed WGS on 32 of these humanized yeasts.

**Figure 2.**
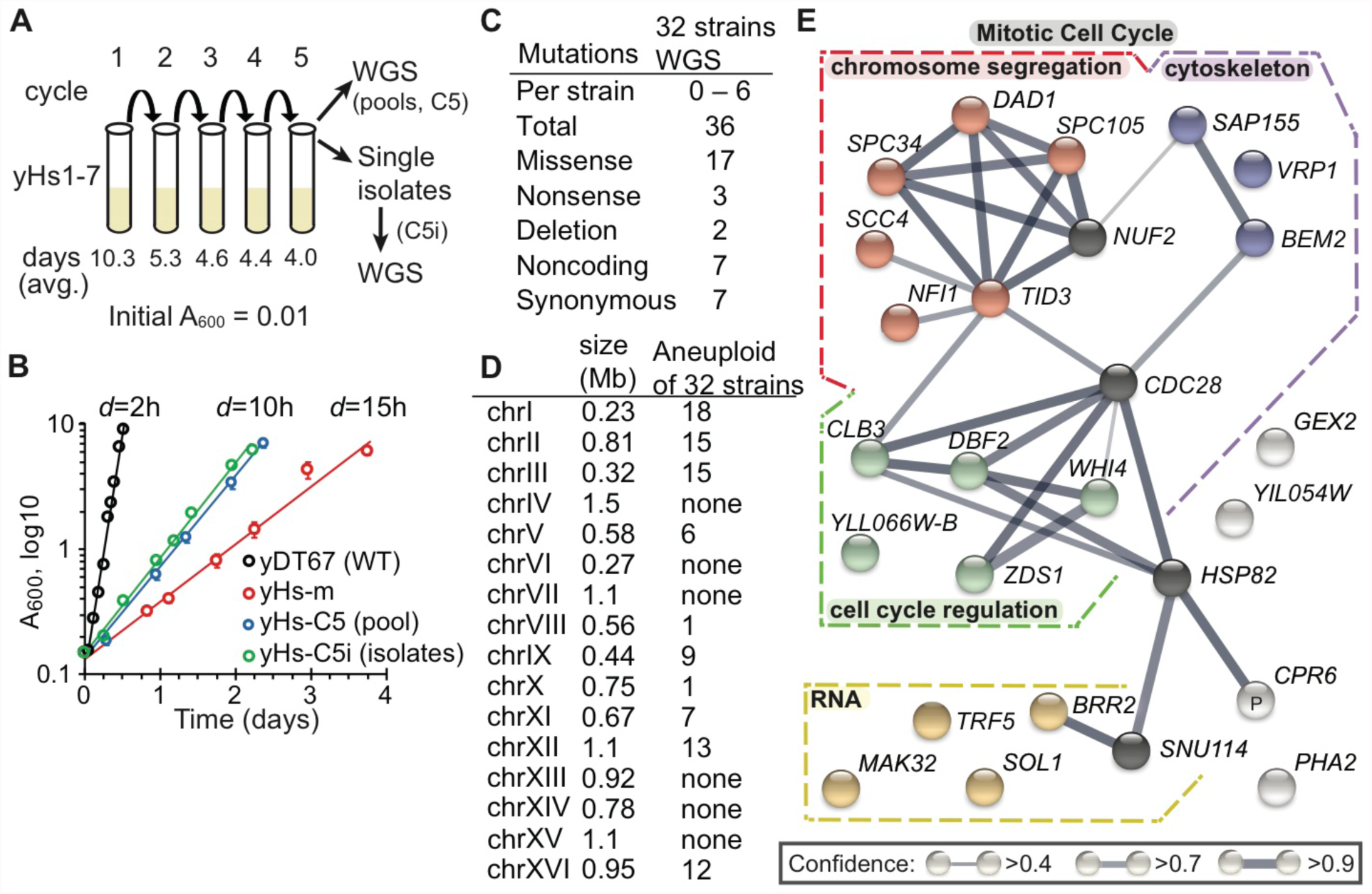
Acquisition of bypass mutations in cell-cycle genes promotes growth with human nucleosomes. (A) Seven yHs-strains were evolved for 5 cycles in liquid medium (SC–Trp). (B) Growth rates and doubling times (*d*) in SC–Trp. Complete doubling times are listed in Table S2. (C) Types of DNA mutations identified by whole-genome sequencing (WGS). (D) Number of times each listed chromosome was scored as aneuploid. (E) 22 unique mutations identified by WGS (Tables S1 and S2) were constructed into a network (*p*=5x10^-5^) using the *String* algorithm (Szklarczyk et al., 2015). Colored nodes are in similar processes. Black nodes are top 4 interacting genes inferred from the network, but not arising as suppressors.

Pulsed-field gel electrophoresis of whole chromosomes from each yHs-lineage showed normal chromosome size (Figure S2D). However, WGS revealed recurrent aneuploidy of specific chromosomes (Figures 2D, S2E, and Table S2). The human-histone plasmid copy number was no more than 2-fold higher than that in the parental strain yDT51. The majority of aneuploidies may be a detrimental consequence of human nucleosomes – as aneuploidies typically reduce fitness (Sheltzer et al., 2011) – but our frequently recurring aneuploidies (Figure 2D) were consistent with other studies (Pavelka et al., 2010), which consider them possible reservoirs for positive selection. Chromosome number was often unstable during lineage evolution, as fractional differences (e.g., 1.5- fold chr1) were not due to insertions/deletions or diploidy, indicating potentially variable levels of aneuploidy at the population level. Only the evolved isolates yHs4C5i1 and the yHs7C5 lineage had a normal chromosomal sequence coverage, the latter perhaps due to a mutation in the gene *DAD1*, which controls microtubule force at the centromere (Sanchez-Perez et al., 2005). By contrast, yHs5 and its progeny had higher levels of aneuploidy, perhaps due to a subtle mutation in the gene *SCC4*, a cohesin loader (Lopez-Serra et al., 2014).

All humanized strains were either missing segments of mitochondrial DNA (mtDNA)(ρ^-^) or showed complete loss of mtDNA (ρ^0^), except for the lineages from yHs5 (ρ^+^). We investigated whether mtDNA loss alone might explain the slow growth rates (Veatch et al., 2009), but found that isogenic-WT ρ0 cells grow better than all humanized lines, and moreover the ρ^+^ yHs5C5i1 isolate was not the fastest growing isolate (Table S2).

We identified 36 mutations in or near genes among the 8 isolates and their derivatives (Figure 1C, Tables S1 and S2), and 22 unique mutations appeared likely to affect gene function based on alterations to protein sequences. We constructed an interaction network from these 22 mutations using the *String* algorithm (Szklarczyk et al., 2015) (Figure 2E). The enrichment of GO terms in this network was non-random, as the genes clustered in 4 processes: chromosome segregation, cytoskeleton, cell-cycle progression, and genes affecting RNA metabolism. These first 3 processes collectively affect mitotic cell-division (Janke et al., 2001; Lew and Reed, 1995). Therefore, mutations in genes that affect cell-division may suppress defects arising from human histones, possibly by circumventing cellular checkpoints triggered by aberrant chromatin properties. These results illustrate how much easier it is to evolve the surrounding the gene network to accommodate new functions rather than the gene itself.

### Specific residues in termini of human histones H3 and H2A limit yeast growth

We were surprised to not identify any mutations within the human histone genes themselves. Converting the C-terminal residues of human histone H3 back to the yeast sequence enhances complementation (McBurney et al., 2016), and some species-specific residues cause lethality when mutated to alanine (Dai et al., 2008; Nakanishi et al., 2008). We systematically swapped residues from human to yeast across histones H3, H4, and H2A, in order to identify species-specific regions (Figures S3 and S4), but did not perform such studies on H2B as it complemented relatively well.

Swapping-back three residues in the C-terminus of hH3 enhances the humanization frequency (Figure S3A), consistent with a recent study (McBurney et al., 2016), whereas swapping-back the lethal residues provided no benefit (Dai et al., 2008). Although hH4 already worked well, we identified two residues in its C-terminus that enhanced humanization (Figure S3B). Only two swapped-back residues in hH3 (hH3_KK_; human–>yeast P121K and Q125K) were required for complementation when combined with completely human H4 (Figure S3C), although there appear to be differences between hH3.1 versus hH3.3 in this regard. A possible explanation for the two hH3 swap-back residues may be that in yeast H3, the two lysine swap-back residues are ubiquitylated by Rtt101^Mms1^, and mutations in H3 of K121R/K125R reduced H3/H4 dimer release from Asf1, restricting transfer to other histone chaperones (Han et al., 2013).

Swapping-back three broad regions in hH2A enhanced complementation in combination with fully human hH2B, the N-terminus, the C-terminus, and a region from residues 19 to 42 (Figure S4). Further analyses narrowed the essential residues to three residues each in the N-terminus and C-terminus. Combining all six of these residues significantly enhanced the humanization frequency and growth rate of the yeast (Figure S4D). Intriguingly, the mammalian lineage-specific N-terminal arginine residues, when inserted into yeast H2A, have been shown to increase chromosome compaction (Macadangdang et al., 2014). The C-terminal portion, which is exposed on the nucleosome face (White et al., 2001), may interact with histone chaperones (e.g., NAP1) analogous to the H3/H4 interaction with Asf1.

We combined the 3 terminal-regions (hH3.1_KK_, hH2A_N_, and hH2A_C_) into human nucleosomes as various “Swapback strains” (Figure 3). As expected, combining all 3 swapped-back regions enhanced humanization (8-residue swayback strain yDT98) to 10^-2^ in only 3 days (Figure 3B). However, the swapback strain with only the two C-terminal regions (hH3.1_KK_ and hH2A_C_; 5-residue swap strain yDT97) grew as fast as the 8-residue swapback version (yDT98), and both of these strains grew nearly as fast as our isogenic-WT strain (yDT67) in 3 days. The 5-residue swapback strain (yDT97) was used for further studies.

**Figure 3.**
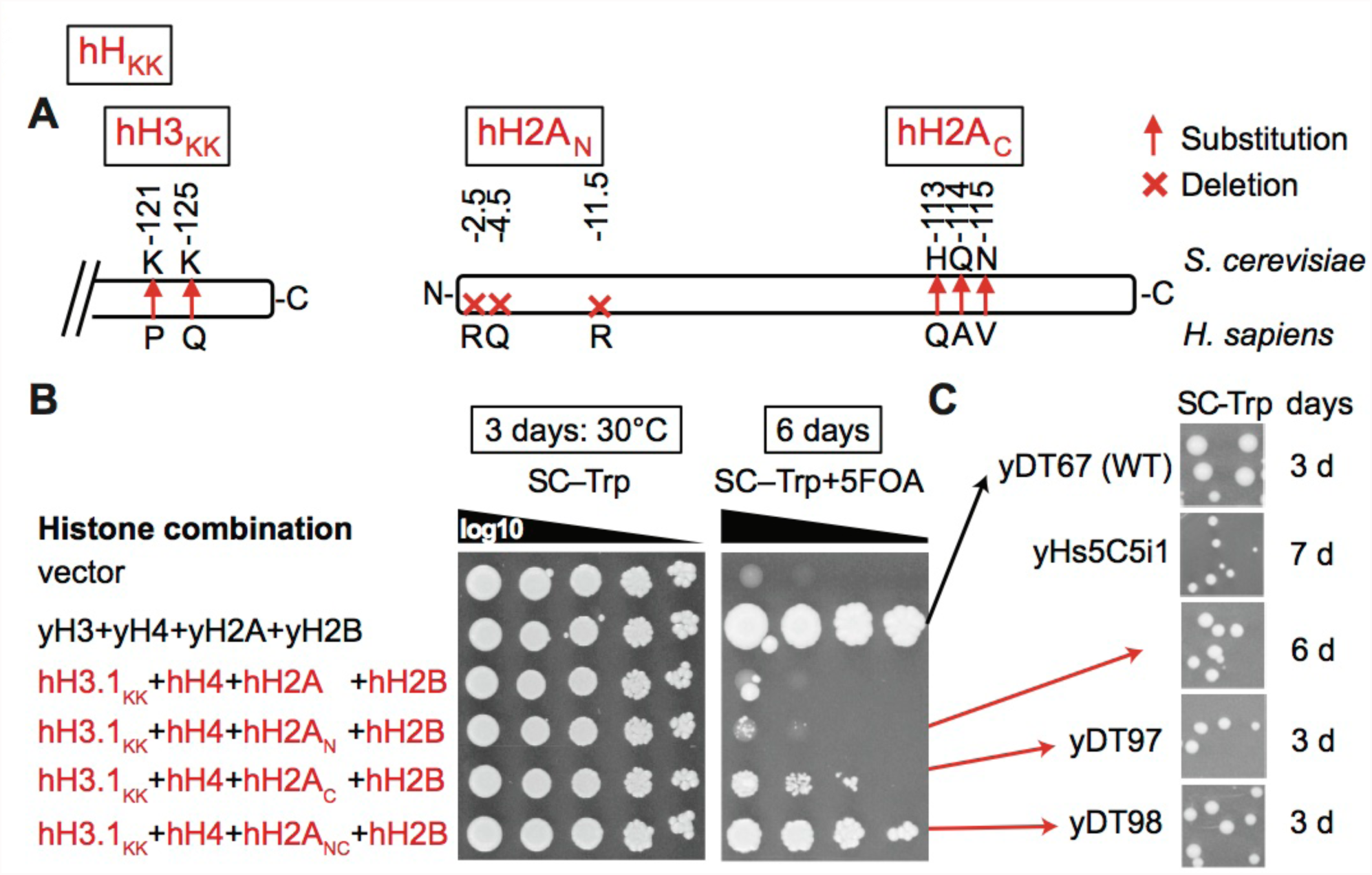
Specific residues in the C-termini of histones H3 and H2A limit growth rates. (A) Maps of swapback residues that enhance human histone utilization identified in Figures S3 and S4. Two residues in the C-terminus of human histone H3 (hH3), and three swap-back residues each in the N-terminus or C-terminus of human histone H2A (hH2A) improved the complementation frequency and growth rate in conjunction with their respective human histone counterpart (i.e., hH4 and hH2B respectively). (B) Systematic combinations of swapback residues in hH3 and hH2A along with hH4 and hH2B show that eight swapback residues promote the highest rates of complementation. (C) Colony growth rate analyses shows that the five-residue swapback strain (yDT97) grows as well as the eight-residue swapback strain (yDT98). Both swapback strains grow at rates closer to isogenic-WT yeast (yDT67), and better than the fastest growing completely humanized isolate (yHs5C5i1).

### Human nucleosomes delay adaptation to new transcriptional programs in yeast

Intriguingly, we often observed that the humanized cells had difficulty adapting to new environments (e.g., colony to liquid culture), which suggested slowed chromatin remodeling to new transcriptional programs. Consistent with this hypothesis, using a *GAL1*-promoter driven eGFP as a proxy for switching to the galactose utilization transcriptome using the RSC complex (Floer et al., 2010) we showed that cells with human nucleosomes had a pronounced delay in transcriptional response to galactose as the sole carbon source, as well as decreased maximal expression on induction (Figure 4A).

**Figure 4.**
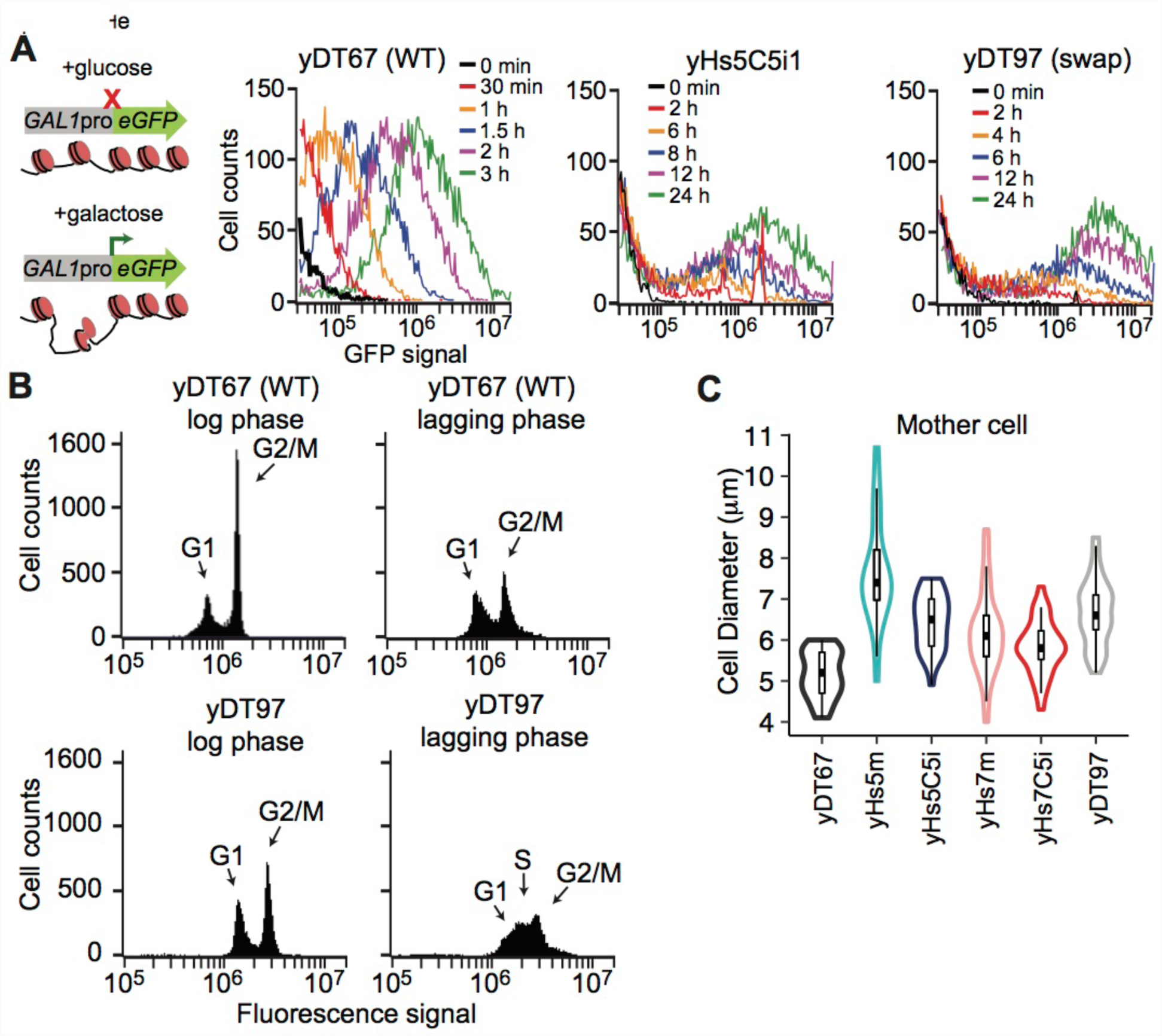
Humanized yeast have trouble adapting to new conditions. (A) Humanized yeast have delayed chromatin remodeling. Chromatin remodeling time-course was analyzed by galactose induction of eGFP using flow cytometry. (B) Humanized yeast have a prolonged S-phase and/or arrest in G1. Cell-cycle analysis based on DNA content. Cells were stained with sytox green, and DNA content was measured by flow cytometry. Each plot shows 10,000 cells in log-phase growth, except where indicated. (C) Violin plots showing that humanized yeast cells are larger and have dysregulated cell size based on phase-contrast microscopy measurements.

We then assessed how readily the cells adjust to new phases of the cell-cycle, a process that also requires extensive chromatin remodeling. Using both bud-counting and flow cytometry of log-phase cells, we observed reduced cell-division, as only 40-60% of humanized cells reach the G2/M phase compared to ~90% in isogenic-WT (Figure S5B). More importantly, the lag-phase cultures of the yDT97 “swap” strain display a prolonged S phase, indicative of a delay in adjusting to log-phase growth (Figures 4B and S5D). This could result from an inability to accumulate new histones onto nascent DNA or an inability to remodel and remove chromatin-bound factors (Ma et al., 2015).

The humanized cells were also larger in size on average and produced a greater range in cell sizes (Figures 4C and S5), which could indicate an inability to regulate cell-size control due to less permissive chromatin. By micro-manipulating single cells onto YPD plates, we found no growth difference between large and small cells (Figure S5E). However, unbudded cells (G1) were less likely to continue to grow than budding cells, although they all mostly remained intact after several days of monitoring. Surprisingly, a high fraction of single cells grew for a number of cell-divisions before arresting as a population (i.e., arrested before reaching the size of a visible colony). Together, these results are consistent with the hypothesis that human nucleosomes delay adaptation to other phases of the cell-cycle.

### Nucleosome organization is specified by the chromatin remodeling network

To evaluate the organization of human nucleosomes on the yeast genome we performed MNase digestion titrations and MNase-seq on evolved and swapback strains using ‘high’ and ‘low’ enzyme concentrations (Figures 5 and S6), to reveal possible differences in nucleosome accessibility (Kubik et al., 2015).

**Figure 5.**
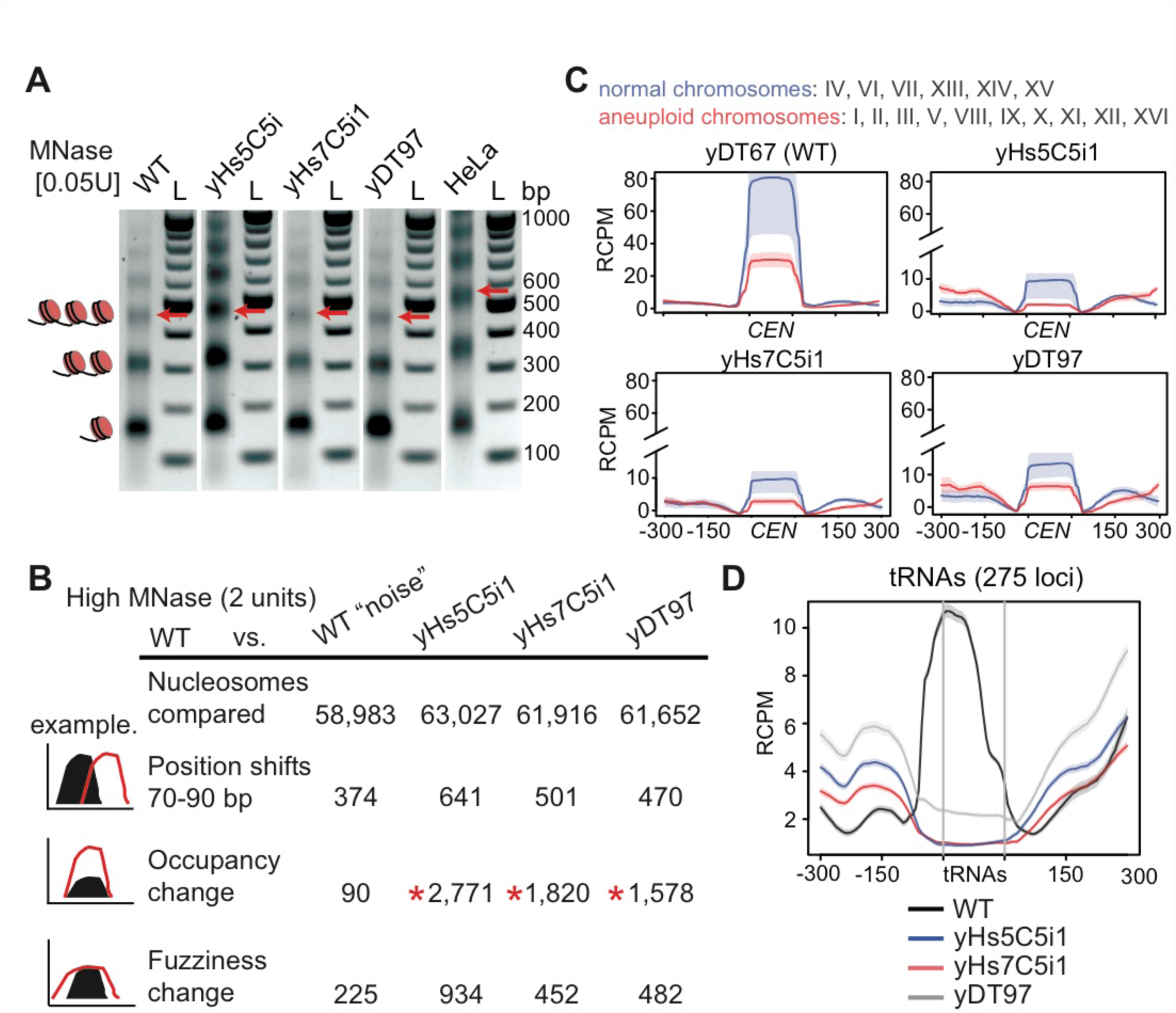
Human nucleosome organization in yeast. (A) MNase digestions reveal that human nucleosomes produce the same nucleosome repeat length as yeast nucleosomes, compared to the longer length of human nucleosomes in HeLa cells. Red arrows indicate position of the tri-nucleosome. The “bp” indicates base-pair size of the DNA ladder (“L”). (B) Table of high (2 units/ml) MNase-seq nucleosome dynamics between humanized to WT yeast, and WT experiment 1 to WT experiment 2 (“noise”). Occupancy and fuzziness changes use a strict False Discovery Rate cut-off of 0.05 (p < 10^-85^) and additional parameters in *Methods*. (C) High MNase-seq read counts at centromeric regions, plotted for chromosomes that were normal or aneuploid in Figure 2D. RCPM refers to read counts per million mapped reads. (D) High MNase-seq read counts for all 275 tRNA genes comparing humanized vs. WT strains showing depletion of either RNAP3 or nucleosomes.

Unexpectedly, the nucleosome repeat length (NRL) of yeast chromatin built using human nucleosomes was identical to the NRL in isogenic-WT yeast, and is substantially shorter than that for human HeLa cells (Figure 5A). The dinucleosome length (~300 bp) from low concentration MNaseseq confirms a short mean nucleosome repeat length (Figure S6C). These data indicate that the NRL in humans is not an intrinsic property of human core nucleosomes, but is likely specified by nucleosome remodelers, by the genomic sequence itself (Segal and Widom, 2009) or by some combination of these factors.

To our surprise, the numbers of nucleosomes with altered positioning or fuzziness (movement) was no different than that of isogenic-WT “noise” (Chen et al., 2013). However, there are substantial occupancy differences, which are distinct even amongst the humanized lines (Figure 5B). Nevertheless, this suggests that nucleosome positioning is determined less so by the type of nucleosome, and much more so by the underlying DNA sequences and the network of chromatin-remodelers for a given species.

As suggested by the chromosome segregation suppressor mutations identified earlier (Figure 2E), we find that human nucleosomes lead to depletion of centromeric nucleosomes as well as relative to the surrounding nucleosomes, perhaps due to conflict with the yeast centromeric H3 variant *CSE4* (Figures 5C and S6D). Relative to the neighboring nucleosomes, depletion was greatest for centromeres on aneuploid chromosomes observed earlier by WGS (Figures 2D and 5C). Strain yHs5C5i1, which had the highest levels of aneuploidy, had greater depletion at these nucleosomes, whereas strain yHs7C5i1, which has normal chromosome numbers and carries a relatively subtle missense mutation (E50D) in the essential gene *DAD1*, has slightly better positioning at these nucleosomes (Figure 5C).

Finally, all 275 tRNA genes had depleted sequence coverage in their gene-bodies compared to WT (Figure 5D). Unlike RNAP2 genes, tRNAs possess an ‘internal control region’, thus, the depleted regions could represent a loss of RNAP3 and accessory factors (Acker et al., 2013), or nucleosome depletion coupled to RNAP3 transcription elongation. In fact, substantially elevated tRNA levels were observed in RNA from yHs cells (Figure S7A), perhaps suggesting human nucleosomes are less stably bound to tRNA sequences. However, as yeast tRNAs are already highly expressed and mostly devoid of nucleosomes, this could instead indicate that tRNA levels are normal, and that it is mRNAs that are highly repressed by human nucleosomes, thus altering the tRNA/mRNA ratio.

### Human nucleosomes produce chromatin more generally repressive for RNAP2

As predicted, total RNA content – predominantly mRNA and rRNA – is reduced by 6-8 fold in all the pre-evolved humanized yeast, and only slightly increased in the evolved and swapback strains (Figure 6A). The mRNA to rRNA ratios remain similar to our isogenic-WT strain (Figure S7A). However, tRNA sized molecule(s) are elevated relative to total RNA, and this may alter the balance of RNA types in the cell. However, we found the reduced total RNA is not explained by substantially altered cell numbers per A_600_ or reduced cell viability as determined by sytox green or trypan blue staining of dead cells (Kwolek-Mirek and Zadrag-Tecza, 2014) (Figure S7B, C). Humanized whole-cell extracts had similar bulk protein yields to isogenic-WT, but the SDS-page gel stained with Coomassie shows numerous proteins with reduced levels, consistent with reduced RNA (Figure S7D), whereas other presumably highly stable proteins are relatively unaffected. Immunoblots using antibodies more specific for human H3 and H4 show greater signal for humanized strains (Figure 6B). Finally, both H3K4 trimethylation and H3K36 trimethylation signals were similar to the isogenic-WT strain, as these modifications are in regions conserved between yeast and humans. This suggests that low mRNA levels are not due to changes in these histone modifications.

**Figure 6.**
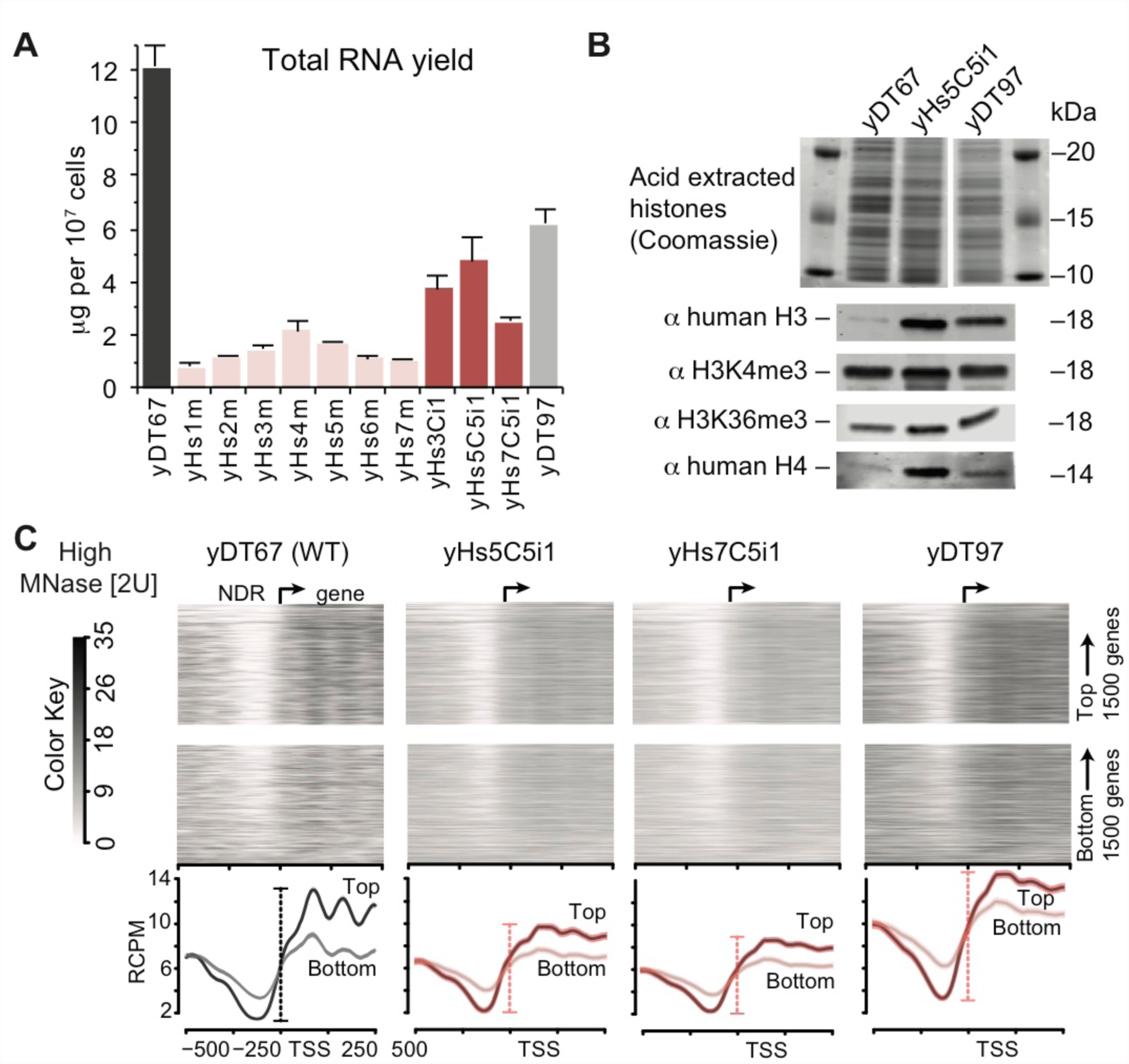
Human nucleosomes are more repressive. (A) Pre-evolved yHs strains (yHs-m) have reduced levels of bulk total RNA (6-8 fold), whereas the evolved and swap strains slightly increase RNA content. Bars show standard error of the mean of 3 biological replicates. (B) Acid-extracted histones from strains analyzed for equal loading by Coomassie staining, and then immunoblotted using different H3 and H4 antibodies. (C) Heatmaps and average profiles of high concentration MNase-seq reads aligned around the transcription start sites (TSS) ±500 bp of the top and bottom 1500 genes by expression. RCPM and color key refers to read counts per million mapped reads.

The reduced RNA content and slowed growth might reflect differences in nucleosome dynamics (Chen et al., 2013), and could indicate a fundamental property of human histones or their relative inability to interact with yeast chromatin remodelers. To understand this effect, we mapped the MNaseseq reads across the transcription start sites (TSS) of the top 1500 genes by expression, and the bottom 1500 genes by expression. Genome-wide, the MNase-seq reveals a less “open” nucleosome depleted region (NDR) upstream of the TSS for humanized yeast than that found in isogenic-WT yeast (Figures 6C and S7E). The amplitude (difference between the NDR and gene bodies) is smaller for fully humanized yeast compared to isogenic-WT, suggesting reduced RNAP2 transcription that is consistent with the decreased total mRNA/rRNA content (Figure 6A). The amplitude of highly expressed genes in humanized yeast looks more similar to the amplitude of the lowest expression genes in isogenic-WT. Even amongst themselves, the humanized strains show distinct nucleosome profiles, suggesting chromatin heterogeneity at the population-level. Furthermore, nucleosome fragment lengths at high concentration MNase of isogenic-WT yeast show a greater fraction of sub-nucleosomal particles (90-125 bp) compared to humanized yeast (~147 bp), suggesting that human chromatin is less accessible to MNase (Figure S6C).

The above results combined with the poor environmental adaptability and cell cycle delays suggest that human nucleosomes are more generally repressive to RNAP2 transcription than yeast nucleosomes, possibly because they have a higher intrinsic affinity for DNA (the model we favor), thus making them more static, or are less easily removed by the yeast chromatin-remodelers, which did not coevolve with these histone sequences. Such a finding is consistent with the relatively more unstable nature of yeast chromatin *in vitro* compared to human chromatin (Leung et al., 2016), as well as their biology – yeast genes are predominantly expressed (Rando and Winston, 2012) – while humans repress the majority of the genome in virtually all cell types (Djebali et al., 2012; Talbert and Henikoff, 2017; Thurman et al., 2012) (Buschbeck and Hake, 2017). Thus, specialized histone variants with higher DNA affinity and stronger gene repression, might enable multicellular organisms to generate a larger variety of transcriptional landscapes.

### Suppressor mutations and human chromatin “memory” enhance humanization frequency

The numerous suppressor mutations identified earlier (Figure 2) may counteract the various defects observed in yeast with human nucleosomes. If the suppressors make human histones more tolerable, they would be predicted to enhance the “humanization frequency”. To determine this, we re-introduced the native yeast histone plasmid into 13 humanized suppressor strains (Figure 7A; black dots), and allowed mitotic loss of the human histone plasmid, thus reverting these cells to native yeast chromatin, whereupon their growth properties improve. These lines were used for dual-plasmid histone shuffling as before, by re-introducing the human histone plasmid. The humanization frequencies, 10-100 fold greater than for non-suppressor ρ^0^ or ρ^+^ strains, confirms that the identified suppressors enhance tolerance of human nucleosomes. Furthermore, humanized colonies appeared as early as 12 days instead of the 20 originally required. These frequencies are high enough to support histone shuffling using any histone variant or from potentially any species.

**Figure 7.**
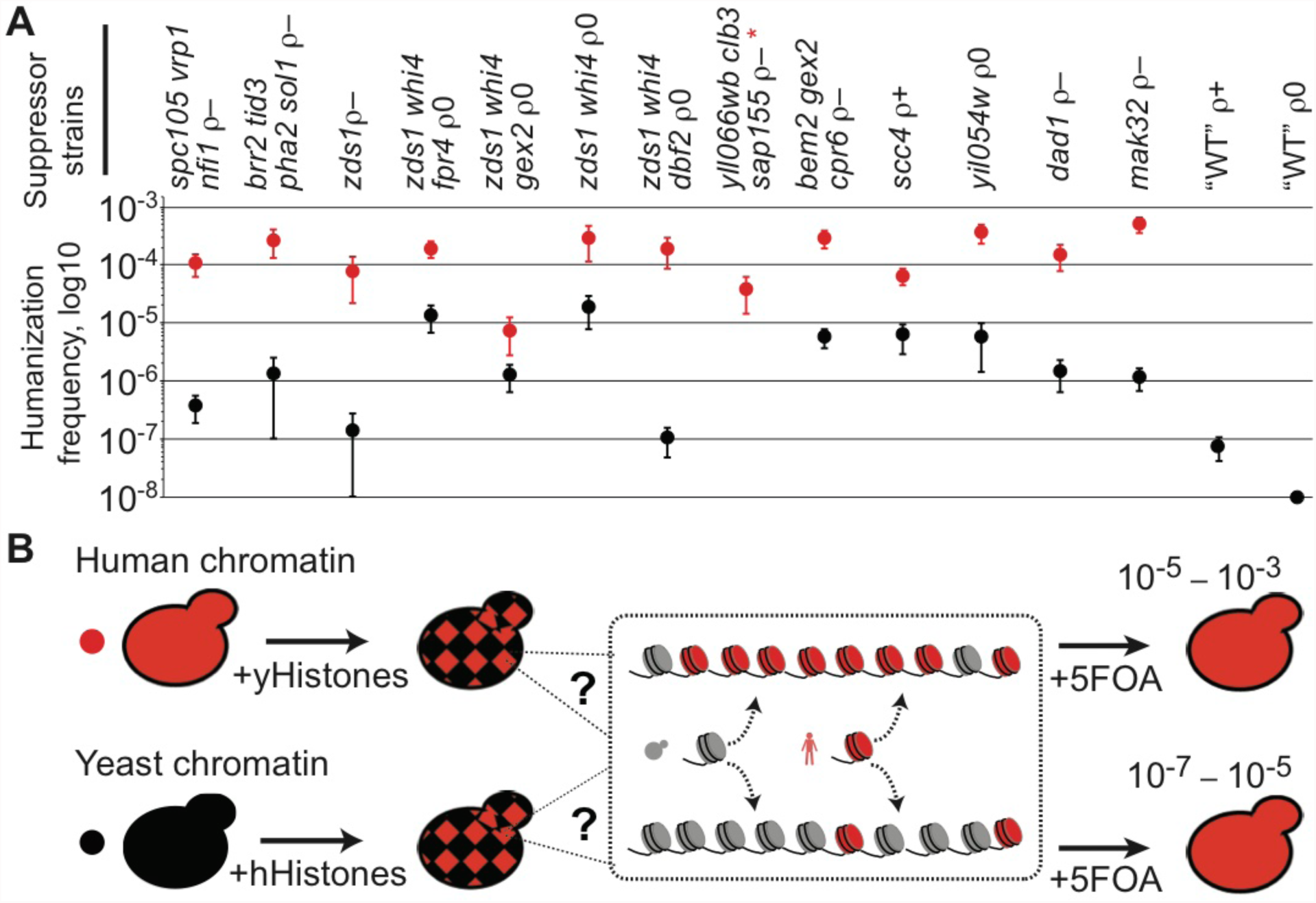
“Re-humanization” of suppressor mutants starting with human or yeast chromatin. (A) Suppressor strains were “re-humanized” (*see* (B)), by generating a mixed chromatin environment (yeast + human), either starting from human chromatin (red dots) or native yeast chromatin (black dots). Genotypes are listed at top. The yHs suppressor strains were converted to a mixed chromatin environment, and then assayed by the dual-plasmid shuffle experiment. Each dot represents the mean of 3 or more experiments on a log10 scale, and the bars are standard error of the mean. Suppressor strain with red asterisk was unable to lose human histone plasmid. (B) Diagram of suppressor strain “re-humanization” frequency experiments. Partitioning of nucleosomes between cells is poorly understood, especially in the context of variant histones. The dotted line inset with a question mark shows the possible chromatin states that may apply when yeast and human nucleosomes coexist and partition to new cells, showing how they may prefer their own histone sub-types.

While the suppressor mutations improved the humanization frequency going from native yeast chromatin to fully human chromatin, we also contemplated how readily human chromatin resists “invasion” by native yeast histones. If the humanization frequency improves in this scenario relative to the above, this might suggest maintenance of human chromatin, and a preference for re-incorporation of nucleosomes of their own type. To test this hypothesis, we reintroduced the native yeast histones into the fully humanized suppressor strains, and allowed for single colonies containing both types of chromatin to grow for approximately 26 cell division generations (Figure 7A; red dots). This is a suitable time frame for native yeast chromatin to outcompete human chromatin, if not for “parental” nucleosome maintenance. Indeed, upon performing 5-FOA plasmid shuffling to remove the native yeast histones, the humanization frequencies reached 10^-5^-10^-3^, and cells appeared as early as 7 days.

We interpret this result to suggest that pre-existing human chromatin might help maintain chromatin of its own kind, at least regionally – a type of chromatin “memory” and transgenerational inheritance – thus pre-disposing some small fraction of cells to resist native yeast chromatin even after many cellular divisions. These results are surprising, as with few exceptions yeast do not have protein machinery dedicated to maintenance of different histone variants, let alone for human histones. In our model, (Figure 7B) nucleosomes prefer their own type, thus seeding and maintaining similar chromatin domains. Therefore, different histone-types or nucleosome compositions are less likely to invade and outcompete this “parental” chromatin during the partitioning of chromatin during cellular division. This may occur in all cells or perhaps a smaller fraction of “older” cells that retain more of the earlier human chromatin. Finally, these results are consistent with our earlier observation demonstrating the relatively static nature of human core nucleosomes, thus making them more likely to maintain a certain epigenetic state.

## Discussion

Because histones are some of the most conserved genes amongst eukaryotes, it was surprising that fully human nucleosomes so rarely led to *bona fide* humanized yeast. This speaks to the centrality of nucleosomes in regulating diverse cellular processes, including transcription and chromosome structure and movement. Cumulatively, our data suggest that human histones in yeast are deposited less efficiently, possibly due in part to sequence incompatibilities mapping to only 5 residues in the C-termini of H3 and H2A. When they do get deposited, the human histones lead to greater gene repression via less accessible NDRs, variations in chromatin, delayed environmental adaptation resulting from slowed chromatin remodeling, and depleted centromeres that possibly limit kinetochore assembly. The sum of these effect leads to partial cell-cycle arrest in G1 and a slower S-phase, which suppressor mutations in these same pathways alleviate.

The human core nucleosome, consisting of H3.1, H4, H2A.1 and H2B.1, may bind DNA more tightly, as it is predominantly deposited during DNA replication, (Campos et al., 2015) and then can remain in place for years, if not for decades, in a terminally differentiated state (Toyama et al., 2013). Earlier studies on *in vitro* reconstitution of yeast and mammalian nucleosomes suggested that mammalian nucleosomes bind DNA more readily, and that yeast nucleosomes are comparatively unstable (Lee et al., 1982). Given that yeast genes are generally euchromatic, and human genes are heterochromatic, this might indicate an evolutionary basis for histone sequence divergence. Yeast, which must readily adapt to new environments, evolved highly dynamic histones, that retain bifunctional characteristics of histone variants found in humans (Rando and Winston, 2012). For instance, yeast H3 acts as both as an H3.1 (replication-dependent) and H3.3 (replication-independent) variant, while yeast H2A acts as both an H2A.1 and H2A.X (DNA-damage) variant (Eriksson et al., 2012). In contrast, human cells must retain transcriptional states corresponding to cellular type. Thus, specialized histone variants with higher DNA affinity and stronger gene repression, enables multicellular organisms to generate more diverse transcriptional landscapes (Buschbeck and Hake, 2017). Indeed, these yeast with human nucleosomes had great difficulty adapting to new environmental conditions, perhaps due to the fundamentally more static biophysical properties of human core nucleosomes (Leung et al., 2016).

Furthermore, our data suggests that histone sequences do not contribute to the nucleosome repeat length (NRL), as the NRL of yeast with human histones remained yeast-like. In higher eukaryotes, longer linker length is partially attributed to linker histone H1 (Fan et al., 2005; Woodcock et al., 2006), but in yeast, it has been shown that expressing the human H1.2 linker had no effect on the NRL (Panday and Grove, 2016). In humans, the NRL ranges from ~178-205 bp, depending on histone modification state (Valouev et al., 2011), with activation marks having the shortest NRL. Thus, the underlying DNA sequence (Segal and Widom, 2009) and the proteins that interact with histones, such as Isw1a (Krietenstein et al., 2016), are more likely to specify this property.

Converting only five residues, 2 in H3, and 3 in H2A promoted relatively robust utilization of human nucleosomes as “Swapback” strains. In the case of human histone H3, the incompatibility with yeast may be attributed to an absence of two lysines required for ubiquitilation in yeast (Han et al., 2013). Human H2A may also poorly interact with yeast histone-interacting genes. However, this finding is still somewhat surprising as numerous other residues differ from yeast to humans that presumably should have larger roles – including many modified residues. As just one example, histone H3 position 42 is a lysine in yeast, but an arginine in humans (Hyland et al., 2011). Numerous other positions differ (Figure 1A), thus, the relative inability to interact and modify histones at these different sites poses a serious question about the cumulative role of histone modifications during cell growth.

Our study suggests a type of chromatin partitioning “memory”, as yeast with pre-existing human chromatin more readily resisted native yeast histones (Figure 7). Histones are displaced from DNA during transcription, replication, and repair, and then reassembled onto DNA strands (Campos et al., 2014). How cells determine which histone sub-type and modification state must be deposited on the parent and daughter DNA strands in the replication fork remains a continuing question (Lai and Pugh, 2017). Based on our results, the restoration of nucleosomes to the parental strand and inheritance to the daughter strand may occur as a type of “semi-conservative replication” of chromatin, whereby both parent and daughter strands retain a portion of the ancestral nucleosome (human), and then may simply attract similarly composed or modified new nucleosomes. This may be a simple way to retain an epigenetic state. Our results argue against a “conservative” model, wherein daughter strands acquire only fresh nucleosomes. This model predicts that humanized yeast transformed with yeast histones, would react similarly to native yeast transformed with human histones (Figure 7B). However, the results leave open the possibility of a “dispersive” model, which is a mixture of the two models. Nevertheless, humanized yeast may permit more systematic study of this process, coupled with future advances in single cell chromatin-profiling methods.

More generally, humanizing the chromatin of budding yeast provides new avenues to study fundamental properties of nucleosomes. We have explored some of these longstanding questions about histone variants: how they alter the dynamics of the genome at the structural and transcriptional level (Talbert and Henikoff, 2017); how they associate into different compositions of nucleosomes (Bernstein and Hake, 2006); and how they are partitioned and repositioned from cell-to-cell across generations (Budhavarapu et al., 2013; Campos et al., 2014). These questions remain fundamental, as many human cells are reprogrammed and differentiate using histone variants during development and disease (Santenard and Torres-Padilla, 2009; Wen et al., 2014). The number of histone variants found in humans is large, and includes many variants that accumulate during aging and disease. Thus, introducing foreign histones from a distant species into the simple yeast genome will help address these many questions.

## Methods

### Strains, plasmids, and media

All yeast strains used in this study were haploid *MAT***α**, except as indicated in Table S5. Yeast to human complementation studies of histones H3 or H4 alone or in combination, were performed in strain yDT17. Strain yDT17 was generated by replacing the *HHT1*-*HHF1* locus with *NatMX4* by one-step PCR recombination, reintroducing *HHT1*-*HHF1* on a *URA3* containing pRS416 plasmid, and then replacing the *HHT2*-*HHF2* locus with *HygMX4*. Experiments involving H2A or H2B alone or in combination used strain yDT30. Strain yDT30 was generated by replacing the *HTA2*-*HTB2* locus with *HygMX4*, transformation with pRS416-HTA2HTB2, and then deleting the *HTA1*-*HTB1* locus as above with *KanMX4*. Analysis of all four histones was performed in strain yDT51. yDT51 was generated similarly to the above, but contains plasmid yDT83 (pRS416-*HTA2-HTAB2-HHT1-HHF1*). The antibiotic markers (*KanMX4, NatMX4, HygMX4*) and the *His3MX4* cassette used in replacing the four loci were reclaimed by deleting these open reading frames using the CRISPR/Cas9 system (DiCarlo et al., 2013) at the positions indicated with red arrows in Figure S1A.

Human histone genes were codon-optimized for yeast and synthesized by Epoch Biolabs. All cloning was performed by Gibson Assembly (Gibson et al., 2009). Swapback residue (human to yeast) histone variants were generated either by gene synthesis or site-directed mutagenesis. A complete list of available strains and plasmids are in the Supplemental Data.

### Dual-plasmid histone shuffle assay

Shuffle strains (yDT17, yDT30, or yDT51), which already contains a set of yeast histones on a *URA3* plasmid, were transformed by a standard Lithium Acetate protocol with a *TRP1* human histone plasmid, which uses the endogenous promoters/terminators from the other yeast histone set. Colonies were selected for 3 days at 30°C on SC–Ura–Trp plates. Single colonies were picked and grown up overnight at 30°C in 2 ml of SC–Trp. Spot assays (as indicated) were diluted 10-fold from overnight cultures (A_600_ of ~10) and spotted on both SC–Trp and SC–Trp+5FOA plates. Shuffle assays for fully human nucleosomes using strain yDT51, were done as above, except 400 μl of overnight culture were spread onto a 10-cm SC–Trp+5FOA plate (25 ml) and incubated at 30°C for up to 20 days in a sealed Tupper-ware container.

### Plasmid isolation from yeast cells

Cells were harvested from 5 ml SC–Trp overnight culture and re-suspended in 600 μl of water and glass beads. Cells were vortexed for 10 minutes to disrupt cells. Plasmids were then isolated by alkaline lysis using the Zymo Zyppy miniprep kit and eluted in 20 μl of water. 5 μl was used to transform *E. coli,* and isolate pure plasmid.

### PCRtag analysis

Crude genomic DNA was generated using a SDS/Lithium Acetate method (Looke et al., 2011). Comparative PCRtag analysis was performed using 0.5 μl of crude gDNA in a 20 μl GoTaqGreen Hot Start Polymerase reaction (Promega) containing 400 mM of each primer (Table S7). Reactions were run as follows: 95°C/5 min, followed by 35 cycles of (95°C/30 s, 62°C/30 s, 72°C/30 s) followed by a 72°C/2 min extension. A 10 μl aliquot was run on a 1% agarose/TTE gel.

### Pulsed-field gel electrophoresis

Intact chromosomal DNA plugs were prepared as described elsewhere (Hage and Houseley, 2013). Chromosome identity was inferred from the known molecular karyotype of parental cells (yDT51) itself derived from *S288C* that was run on the same gel. Samples were run on a 1.0% agarose gel in 0.5x TBE for 24 h at 14°C on a CHEF apparatus. The voltage was 6 V/cm, at an angle of 120° and 60-120 s switch time ramped over 24 h.

### Mating and sporulation tests

Mating tester lawns (*his1* strains 17/17 *MAT**α*** or 17/14 *MAT**a***) were replica plated to YPD plates. A large amount of humanized strains (*HIS1*) were then smeared onto the replica plate to form rectangles, and then incubated overnight at 30°C. Plates were then replica plated onto synthetic defined (SD) plates and incubated overnight at 30°C. The diploids were sporulated for 7 days as previously described (Dai et al., 2008).

### Microscopy

All yeast were grown to an A_600_ of 0.5-0.9 in SC– Trp liquid media, and imaged under phase-contrast conditions at 100X magnification using a Nikon Eclipse Ti microscope. Cellular diameters were measure from 4 images each, comprising a total of 50 single cells. Violin plots and boxplots were generated using the R-package ggplot2.

### Cell counting and viability

Cells were manually counted using a hemacytometer with Trypan blue vital dye under a microscope. Cell viability was also measured by incubating cells in 1 μM Sytox Green solution in PBS, and counting number of fluorescent cells (dead) by flow cytometry on a BD Accuri C6 flow cytometer. Coulter counting was performed using a Millipore Scepter by diluting log phase cultures 1:100 in PBS, and then taking up cells according to manufacturer’s recommendations. Micromanipulation of single cells was performed using a Singer MSM 400 onto YPD plates.

### Cell-cycle analysis using sytox green

Cell-cycle analysis by DNA content was adapted from (Rosebrock, 2017). Yeast were grown to log phase unless otherwise indicated in SC–Trp. Lag-phase for yDT67 took 45 min, whereas yDT97 took 2 h. Briefly, 10^7^ cells were fixed overnight in 70% EtOH overnight. Cells were incubated in 500 μl 2 mg/ml RNaseA solution for 2 h at 37°C. Then, 25 μl of 20 mg/ml Proteinase K solution was added, and cells incubated for 45 min at 37°C. Cells were washed and then stored in 1 ml 50 mM Tris pH 7.5. 50 μl of cells were re-suspended in 1 ml of 1 μM solution of Sytox Green (Thermofisher), and then 10,000 events were analyzed by flow cytometry on a BD Accuri C6 flow cytometer.

### Flow cytometry of *GAL1*-eGFP induction

Strains as indicated were transformed by standard lithium acetate with plasmid pAV115-GAL-GFP, and selected on SC– Leu+2%glucose plates at 30°C. Single colonies were grown overnight at 30°C in SC–Leu+glucose(2%). Cells were washed once in PBS, and then sub-cultured into SC– Leu+galactose(2%)+raffinose(1%) media and incubated at 30°C. For the times indicated, 25 μl of cells were diluted into 0.2 ml PBS and 10,000 events were analyzed by flow cytometry on a BD Accuri C6 flow cytometer.

### ”Re-humanization” of suppressor mutants starting with human or yeast chromatin

Humanized lineages were re-transformed with native yeast histone plasmid pDT83 (*URA3*) using standard Lithium Acetate transformation, and selected on SC–Ura–Trp plates for 4 days at 30°C. To determine the “human histone memory”, single colonies were grown overnight in 2 ml SC–Trp and directly used in dual-plasmid histone shuffle as described above. To determine the “rehumanization” rate of suppressor mutations, single colonies from the above re-transformed strains were grown in SC–Ura for multiple sub-cultures to allow mitotic loss of the *TRP1* human histone plasmid pDT109. Cells were replica plated onto SC–Ura and SC–Trp to identify those containing only the native yeast histones. These strains were then used for another round of dual-histone plasmid shuffle as described above.

### Protein Analysis and Western Blotting

Whole-cell extracts were generated using a modified protocol from (Zhang et al., 2011). Briefly, 10^8^ log-phase yeast cells were re-suspended in 400 μl 0.15 M NaOH and 0.5 mM dithiothreitol (DTT), and incubated for 10 min on ice. Cells were pelleted at top speed for 10 min at 4°C, and re-suspended in 65 μl lysis buffer (20 mM HEPES pH 7.4, 0.1% Tween20, 2 mM MgCl2, 300 mM NaCl, 0.5 mM DTT, and 1 mM Roche Complete protease inhibitor) and an equal volume of 0.5 mm glass beads. Mixture was vortexed at top speed for 10 min in the cold room. Subsequently, 25 μl of NuPAGE (4X) LDS Sample buffer and 10 μl beta-mercaptoethanol was added, and the mixture was heated at >95°C for 10 min. The debris was pelleted and the supernatant was run on a 12% Bis-Tris SDS acrylamide gel and stained with Coomassie blue.

Acid extracted histones were generated by first resuspending 5 x 10^8^ log-phase yeast in spheroblasting buffer (1.2M Sorbitol, 100 mM potassium phosphate pH 7.5, 1 mM CaCl_2_, and 0.5 mM β-mercaptoethanol) containing Zymolase 40T (40 units/ml) and incubating for 20 min at 37°C. Spheroblasts were gently spun down at 3000 rpm for 3 min and then re-suspended in 1 ml of 0.5 M HCl/10% glycerol with glass beads on ice for 30 min. Cells were vortexed at top speed for 1 min every 5 min and kept on ice. Mixture was spun at 16,000 x g for 10 min and the supernatant was added to 8 volumes of acetone and left at -20°C overnight. The following day, mixture was pelleted for 5 min at 16,000 x g, the solution poured off and the pellet was air-dryed. Pellet was resuspended in 130 μl water, and then 50 μl NuPAGE (4X) LDS Sample buffer and 20 μl beta-mercaptoethanol was added, and the mixture was heated at >95°C for 10 min. Supernatant was run on a 12% Bis-Tris SDS acrylamide gel and stained with Coomassie blue, or directly used for Western blotting.

Protein samples run on 12% Bis-Tris SDS acrylamide gel were transferred to membrane (Millipore, Immobilon-FL) using the BioRad Trans-Blot Turbo system according to manufacturer’s recommendations. Membranes were blocked for 1.5 h at room temperature in 1:1 Tris-buffered saline (TBS)/Odyssey blocking buffer (LiCor). Blocking buffer was removed and membrane re-suspended in primary buffer overnight at 4°C containing 1:1 TBS + 0.05% Tween-20 (TBST)/Odyssey and the following antibodies used at 1:2,000 dilution: human H3 (abcam ab24834), H3K4me3 (abcam ab1012), H3K36me3 (abcam ab9050), human H4 (abcam ab10158). The following day, membrane was washed 5 times for 5 min each in TBST/Odyssey, re-suspended in secondary antibody buffer TBST/Odyssey/0.01% SDS with 1:20,000 dilution of both IRDye 800 goat anti-mouse and IRDye 680 goat anti-rabbit (LiCor) for 1.5 h at room temperature. Secondary was washed 5 times for 5 min each in TBST/Odyssey and then imaged using dual channels on a LiCor Odyssey CLx machine.

### Growth assay on various types of solid media

Cultures were normalized to an A_600_ of 10 and serially diluted in 10-fold increments in water and plated onto each type of medium. The following drugs and conditions were mixed into YPD + 2% dextrose +2% agar: benomyl (15 μg/ml; microtubule inhibitor), camptothecin (0.5 μg/ml; topoisomerase inhibitor), hydroxyurea (0.2 M; defective DNA replication), NaOH (pH 9.0; vacuole formation defects), HCl (pH 4.0; vacuole formation defects), and methyl methanosulfate (MMS 0.05%; defective DNA repair). Galactose plates were prepared in Synthetic Complete media + 1% raffinose and 2% galactose.

### Whole genome sequencing and data analysis

Genomic DNA was isolated using Norgen Biotek’s Fungal/Yeast Genomic DNA isolation kit, which included a spheroblasting step and bead-beating step. At least 1 μg of genomic DNA was used for Illumina library preparation using the Kapa Truseq library prep, and we routinely multiplexed 30 yeast genomes on a single HiSeq 4000 or 2500 lane.

Paired-end FASTQ files were aligned with the following pipeline. First, adapters, reads shorter than 50 bp, and poor quality reads near ends, were removed using Trimmomatic (Bolger et al., 2014). Data quality was assessed using FastQC. Processed reads were aligned to a custom genome reference (yDT51H.fa) using Burrows Wheeler aligner (BWA) mem algorithm(Li and Durbin, 2010), and Sam files were converted to sorted Bam files using Samtools (Li, 2011). Variants were called using the GATK “best practices” pipeline for Haplotype caller, custom scripts, and manually verified on the IGV viewer. Variants were identical using Samtools “mpileup”. Read counts for each chromosome were determined from WGS Bam files using Bedtools “genome coverage” (Quinlan, 2014). Chromosome copy number was then calculated by generating boxplots in R using ggplot2. Networks for suppressor mutants were generated by uploading genes into the *String* online server (Szklarczyk et al., 2015). GO-terms were identified using the Panther database (Mi et al., 2016). Of the 37 mutations identified (Tables S1 and S2), 6 synonymous mutations were considered “innocuous” based on their similar codon usage.

### MNase-digestions and MNase-sequencing

Experiments were adapted from (Kubik et al., 2015). Biological triplicate yeast colonies were each grown at 30°C to an A_600_ of ~0.9 in 100 ml of SC–Trp media. Cultures were crosslinked with 1% formaldehyde for 15 min at 25°C, and then quenched with 125 mM glycine. Cultures were washed twice in water, and pellets were then stored at –80°C. To perform MNase digestions, cells were first spheroplasted by suspending pellets in 4 ml spheroplasting buffer (1.2M Sorbitol, 100 mM potassium phosphate pH 7.5, 1 mM CaCl_2_, and 0.5 mM β-mercaptoethanol) containing Zymolase 40T (40 units/ml) and incubated for 20 min at 37°C. Spheroblasts were gently washed twice with spheroblasting buffer, and then re-suspended in 1 ml digestion buffer (1M Sorbitol, 50 mM NaCl, 10 mM Tris-HCL (pH 7.4), 5 mM MgCl_2_, 1 mM CaCl_2_, 0.5 mM spermidine, 0.075% NP-40, and 1 mM β-mercaptoethanol). Samples were split into 500 μl aliquots equivalent to 50 ml culture each. To each sample, micrococcal nuclease (Sigma: N5386) was added to a final concentration for high digestion (2 units/ml) or low digestion (0.2 units/ml) or as specified in Figure S6. Digestions proceeded at 37°C for 45 minutes. Reactions were quenched with 16.6 μl of 0.5 M EDTA. Crosslinks were reversed in 0.5% SDS and 0.5 mg/ml proteinase K, by incubating at 37°C for 1h, followed by 65°C for 2 h. Nucleic acid was extracted with phenol/chloroform twice, followed by chloroform. Nucleic acid was precipitated by adding 50 μl Sodium Acetate (3M, pH 5.2), an equal volume of isopropanol, and spinning for 20 min at 16,0000 x g. Pellets were washed once with 70% EtOH, and then resuspended in 50 μl TE buffer containing 6 kUnitz of RNase A, and incubated for 30 min at 37°C. DNA was then purified using a Zymo DNA clean and concentrator, and eluted in 20 μl. MNase digested fragment DNA was measured by Qubit, and assessed on a 1.5% agarose TTE gel. At least 200 ng of DNA (PCR-free or minimal PCR of 2-3 cycles) for each replicate was used to generate a library for paired-end sequencing on an Illumina Hiseq 4000.

### Nucleosome positioning analysis

MNase-seq FASTQ reads were processed using Trimmomatic (Bolger et al., 2014), FastQC, and then aligned to the sacCer3 reference genome using BWA-mem (Li and Durbin, 2010), and then converted to a sorted Bam file using samtools (Li, 2011). Custom bed files corresponding to the top and bottom 1500 genes, centromere regions, and tRNA regions were used to align MNase reads using Ngs.plot (Shen et al., 2014) to regions as specified. Fragment lengths were obtained from Sam files and plotted using ggplot2. Nucleosome dynamics were analyzed using DANPOS2 (Chen et al., 2013). Custom scripts were used to process the data to reduce erroneously called and altered nucleosomes as based on comparing MNase-seq data from WT experiment 1 against WT experiment 2 (“noise”). Nucleosome shifts passed the threshold when both nucleosome comparisons had aligned reads >300 and when shifts were greater than 70 bp. Nucleosome occupancies required that at least one nucleosome comparison have an aligned read count >300, and the False Discovery Rate (FDR) was lower than 0.05 with a p < 10^-85^. Fuzzy nucleosomes required that both nucleosome comparisons have read counts >300 and an FDR of <0.05.

## Acknowledgements

We thank Adriana Heguy and her staff at the Genome Technology Center at NYU Langone Medical Center for performing high-throughput sequencing. We thank Junbiao Dai for constructing some of the original human histone plasmids and yeast strains used in these studies, and Michael Shen for the *GAL1*pro-eGFP plasmid. We thank Leslie Mitchell, Liam Holt, and Karim-Jean Armache for comments on the manuscript. This work was supported by National Institute of General Medical Sciences of the NIH fellowship F32GM116411 to D.M.T., and DARPA grant to J.D.B.

## Author Contributions

D.M.T and J.D.B. designed the study, analyzed the data, and wrote the manuscript. D.M.T. performed the experiments.

## Author Information

J.D.B. is a founder and director of Neochromosome Inc. J.D.B. serves as a scientific adviser to Modern Meadow, Inc., Recombinetics Inc., and Sample6 Inc. These arrangements are reviewed and managed by the committee on conflict of interest at NYULMC. The topic of this paper is the subject of a patent application. Correspondence and requests for materials should be addressed to.

## Data Availability

All high-throughput sequencing data has been deposited in the SRA database under Accession PRJNA379735, and listed in Supplemental Tables 8-9.

---

**Figure S1.**
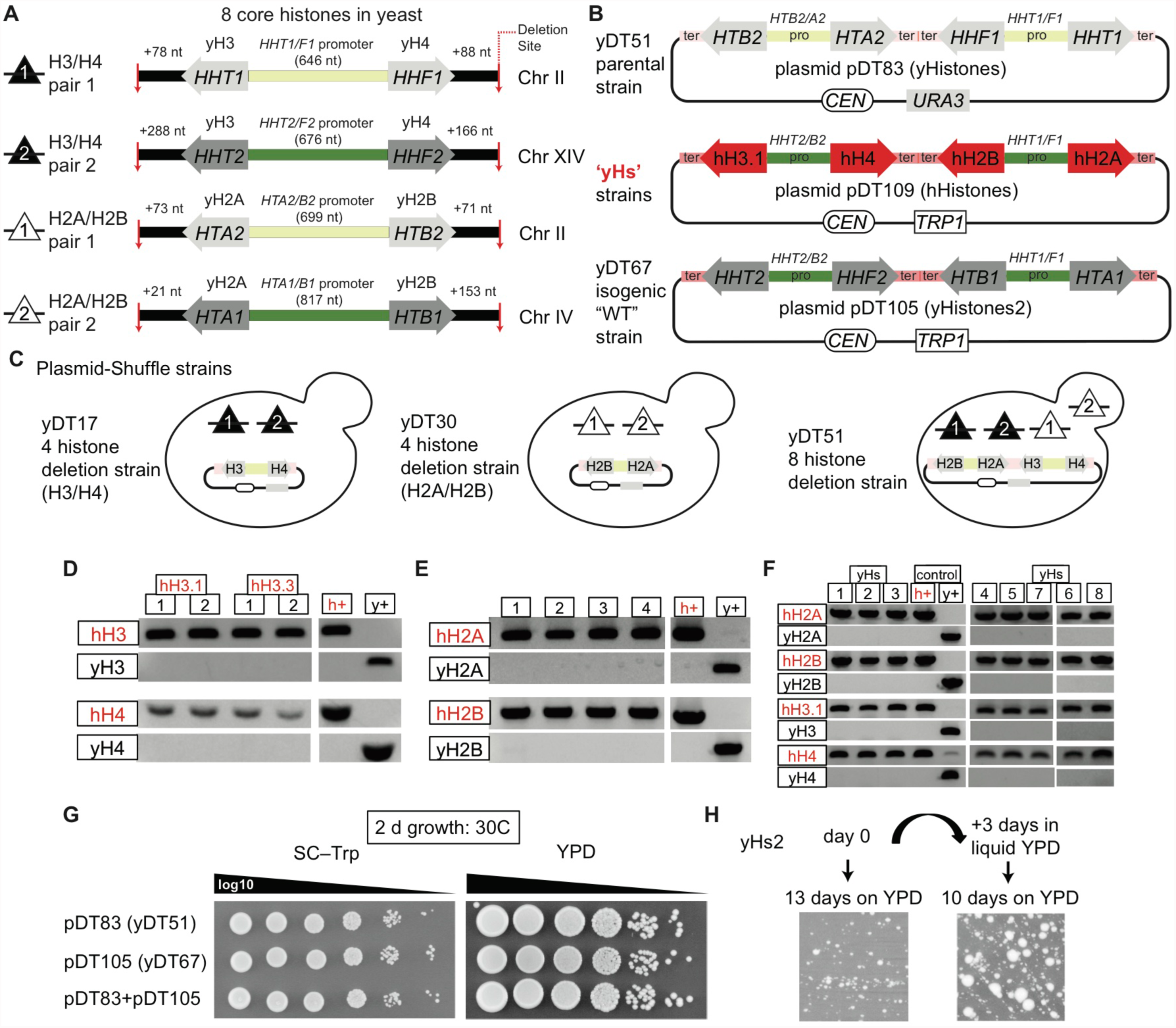
Construction of yeast with completely human nucleosomes (‘yHs’), Related to Figure 1. (A) Map of histone genomic locations in yeast. Triangles show histone pairs deleted in (C). Red arrows indicate CRISPR/Cas9 deletion junctions. Different shades of green show the divergent histone promoters. (B) Diagram of main histone plasmids used in this study for the dual-plasmid histone shuffle. Note the different promoters/terminators used shown in different shades of green. (C) The three histone deletion strains used for humanization studies in Figure 1C, and stabilizing plasmids as indicated. (D) PCRtag confirmation of yeast containing human histones H3.1/H3.3 and H4 (hH3.1/hH3.3 and hH4). (E) PCRtag confirmation of yeast containing human histones H2A and H2B (hH2A and hH2B). (F) PCRtag confirmation of the 8 yeast with completely human nucleosomes with the names “yHs” for “Yeast Homo Sapiens”. (G) Colony growth rates for various “WT” versions of yeast that contain different complements of native yeast histone plasmids. (H) Demonstration of how rapidly “yHs” yeast accumulate suppressors and evolve towards faster growth.

**Figure S2.**
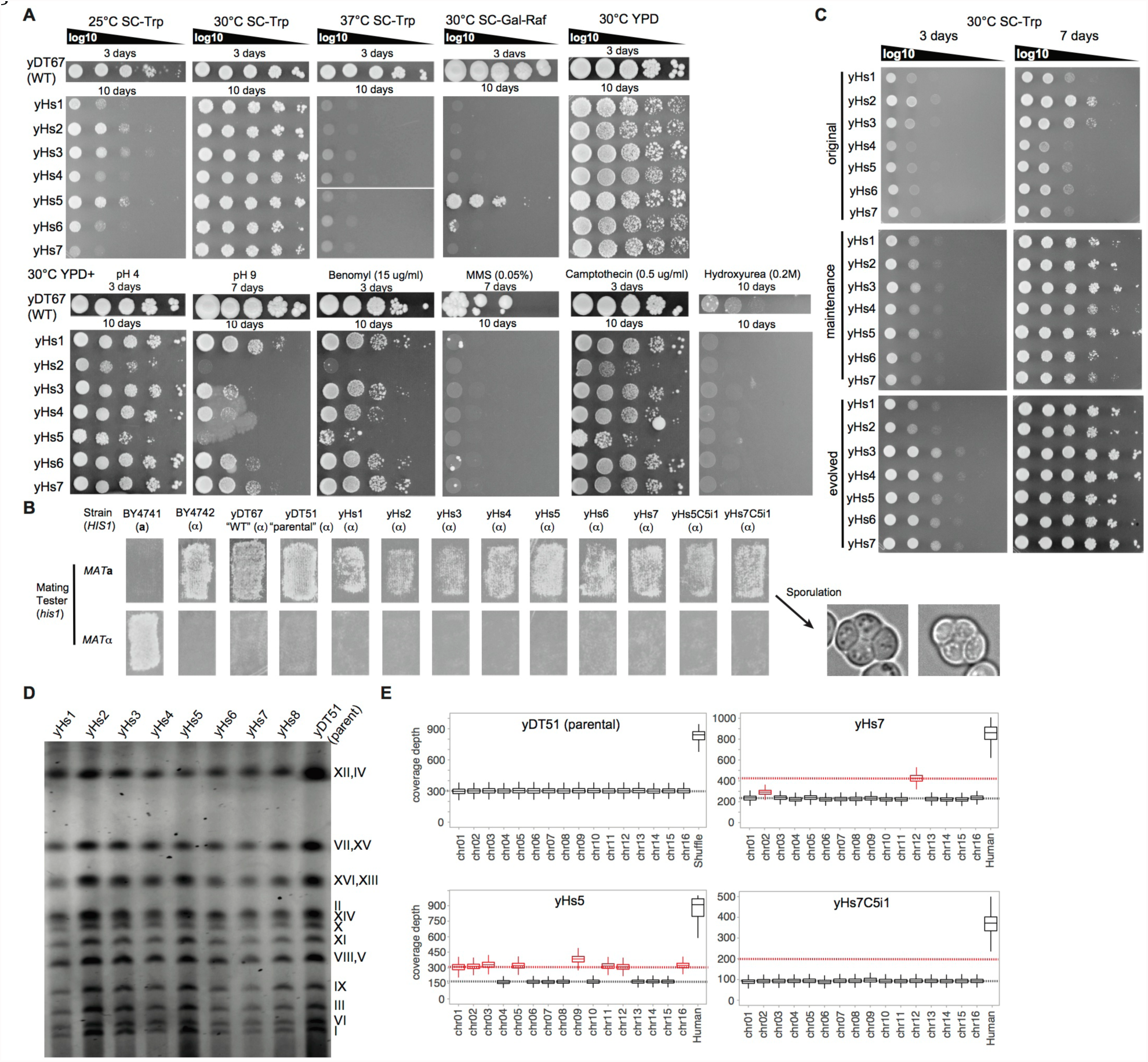
Growth rates of ‘yHs’ strains and chromosomal aneuploidy, Related to Figures 1 and 2. (A) Growth of yHs1-7 on the following drugs and conditions: SC–Trp + 2% dextrose, SC + 1% raffinose and 2% galactose (respiration), YPD + 2% dextrose, YPD + 2% dextrose + either: HCl (pH 4.0; vacuole formation defects), NaOH (pH 9.0; vacuole formation defects), Benomyl (15 μg/ml; microtubule inhibitor), Methyl methanosulfate (MMS 0.05%; defective DNA repair), Camptothecin (0.5 μg/ml; topoisomerase inhibitor), and Hydroxyurea (0.2 M; defective DNA replication). (B) Mating tests of yHs1-7. Mated diploids were then sporulated. (C) Growth comparison of yHs1-7 from original colony isolates, maintenance strains (yHs-m), and evolved strains (yHsC5) on solid media for 3 and 7 days using 10-fold serial dilutions. Cells were normalized to an A_600_ of 10 (D) None of the eight yHs lineages possess gross chromosomal abnormalities (deletions or insertions) as analyzed by pulsed-field gel electrophoresis. (E) Examples of chromosomal aneuploidies for 3 yHs lineages, including yHs7 (aneuploid) and yHs7C5i1, which showed no aneuploidies and acquired a mutation E50D in gene *DAD1*. Box-plots show read counts across each chromosome.

**Figure S3.**
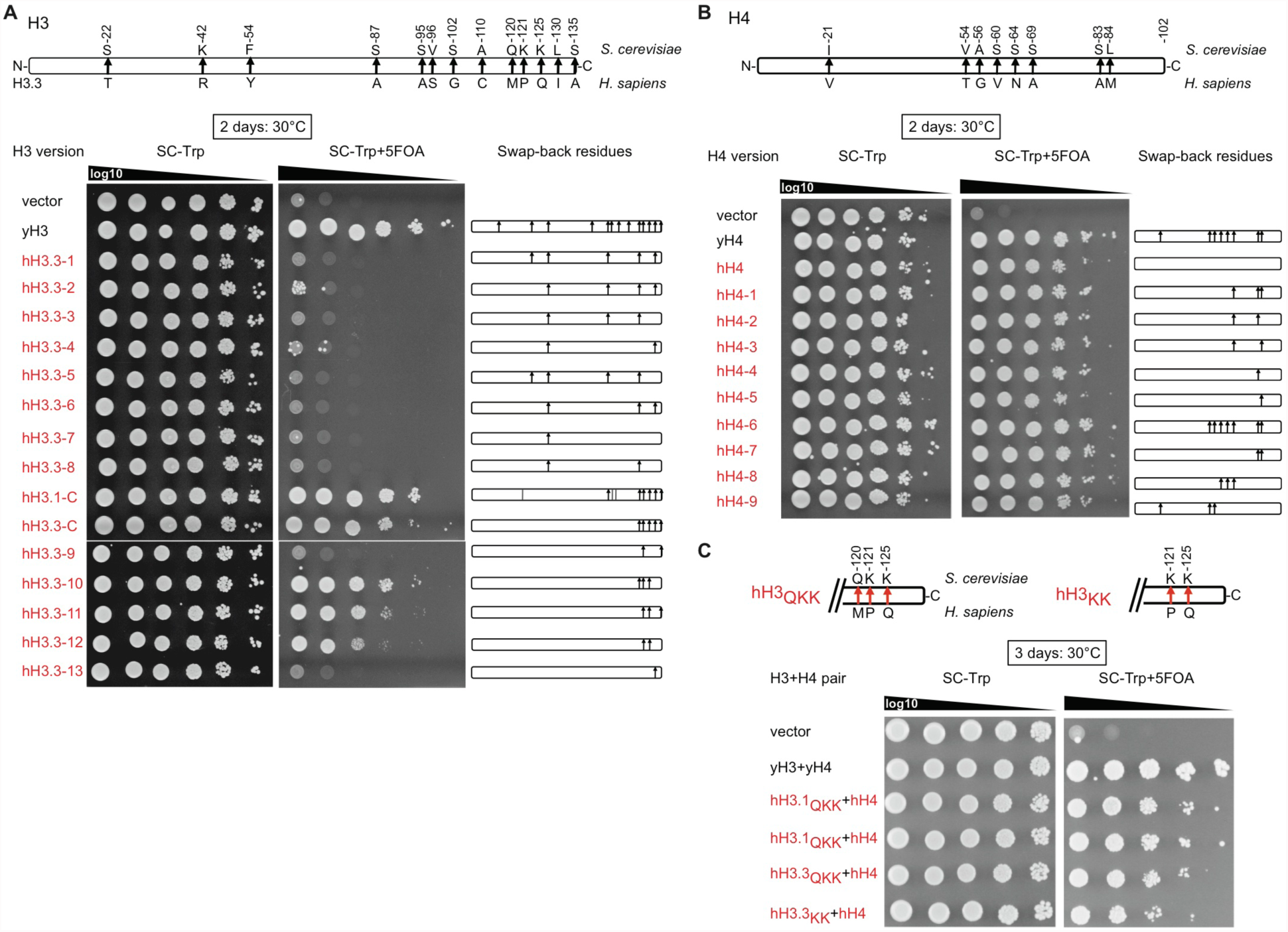
Identification of residues in human H3 and H4, when swapped-back to yeast, improve humanization frequency, Related to Figure 3. (A) Systematic mapping of human to yeast residues in human histone H3.3 using 5-FOA plasmid shuffling. The right-hand shows maps of the tested mutants, with black-arrows indicating positions swapped-back to yeast. Each plasmid was tested in strain yDT17, which contains deletions of both H3/H4 loci and is stabilized with a *URA3*- *CEN* plasmid containing the *HHT1*-*HHF1* locus. Yeast are spotted in 10-fold serial dilutions. Versions labeled hH3.1-C and hH3.3-C were shown to complement well in yeast (McBurney et al., 2016). (B) Systematic mapping of human to yeast residues in histone H4. Swapped-back residues in hH4 were tested as described in (A) also in strain yDT17. (C) Combination of different hH3-swapback strains with completely human H4. When combined with human histone H4 (hH4), only two swap-back residues (P121K and Q125K) are required for hH3.1, whereas three are required for hH3.3.

**Figure S4.**
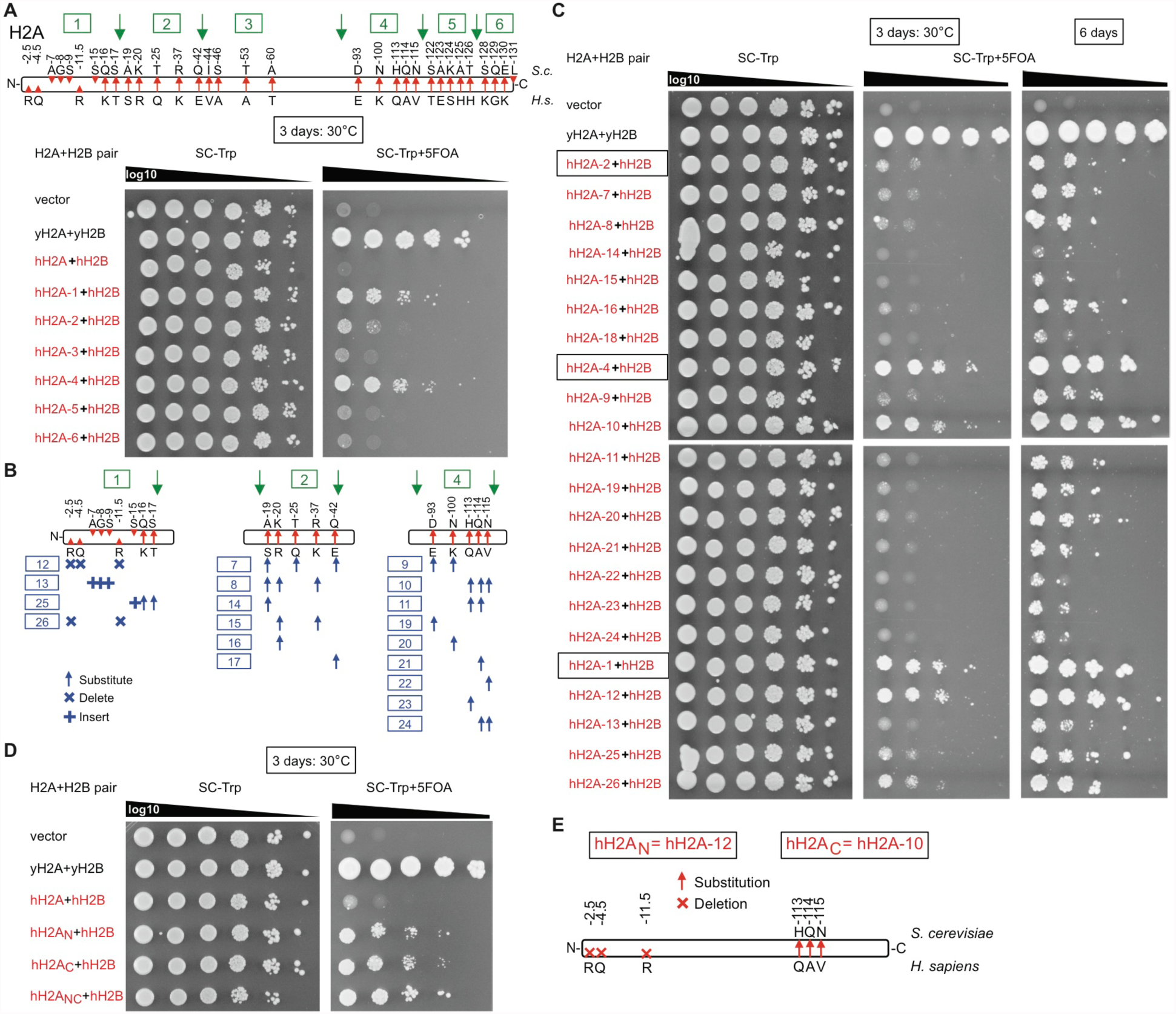
Identification of swap-back residues in human H2A, that improve humanization frequency, Related to Figure 3. (A) hH2A was partitioned into 6 regions, and each region was swapped-back to yeast to test complementation frequency using 5-FOA plasmid shuffling in strain yDT30. (B) Regions 1, 2, and 4 were partitioned into further systematic swap-backs. (C) Complementation assays of swap strains from (B). (D) Three swapped-back residues each in the N-terminus (hH2A_N_) or C-terminus (hH2A_C_) of human histone H2A (hH2A) enhanced humanization frequency and growth rates in combination with human histone H2B (hH2B). The combination of all six swapped-back residues (hH2A_NC_) is optimal.

**Figure S5.**
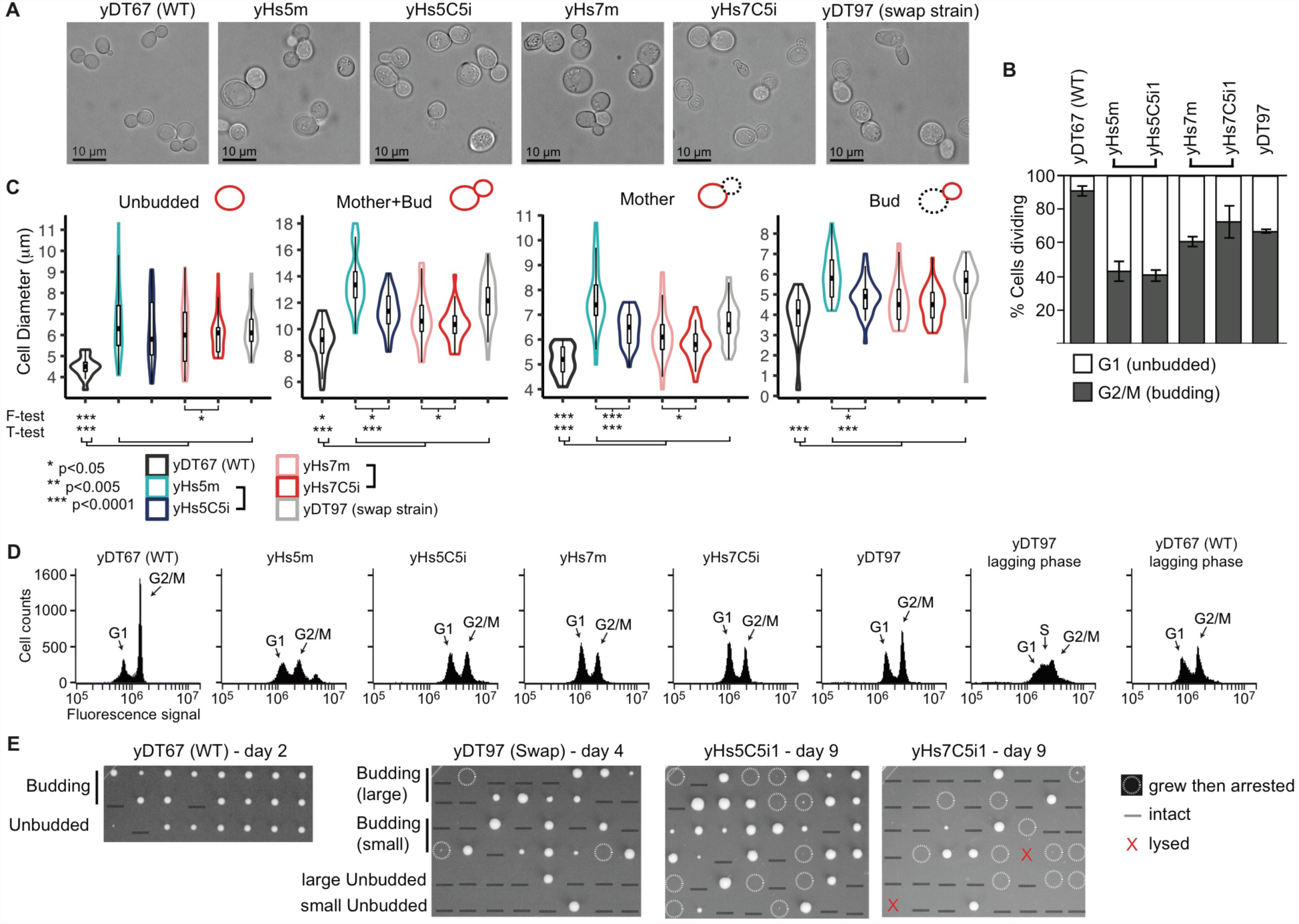
Yeast with human nucleosomes have larger and less regulated cell sizes, and arrest in G1, Related to Figure 4. (A) Image stills of yDT67 (WT) yeast compared to humanized strains. (B) Percent of cells in either the unbudded or budded state from phase-contrast microscopy images. Bars are standard error of the mean from 4 separate images. (C) Violin plots with boxplots inside showing size distributions of the indicated strains for various states of budding. Plots are based on ~50 cells measured from four separate microscopy images. F-tests measure significance of whether two populations have different size distributions. Two-tailed T-tests measure significance of difference in average cell size. (D) Cell-cycle analysis based on DNA content. Cells were stained with sytox green, and DNA content was measured by flow cytometry. Each plot shows 10,000 cells in log-phase growth, except where indicated. (E) Micromanipulation of single-cells for growth. Most cells remained intact (black underline). Cells with white circles grew for a few cell-divisions and then arrested.

**Figure S6.**
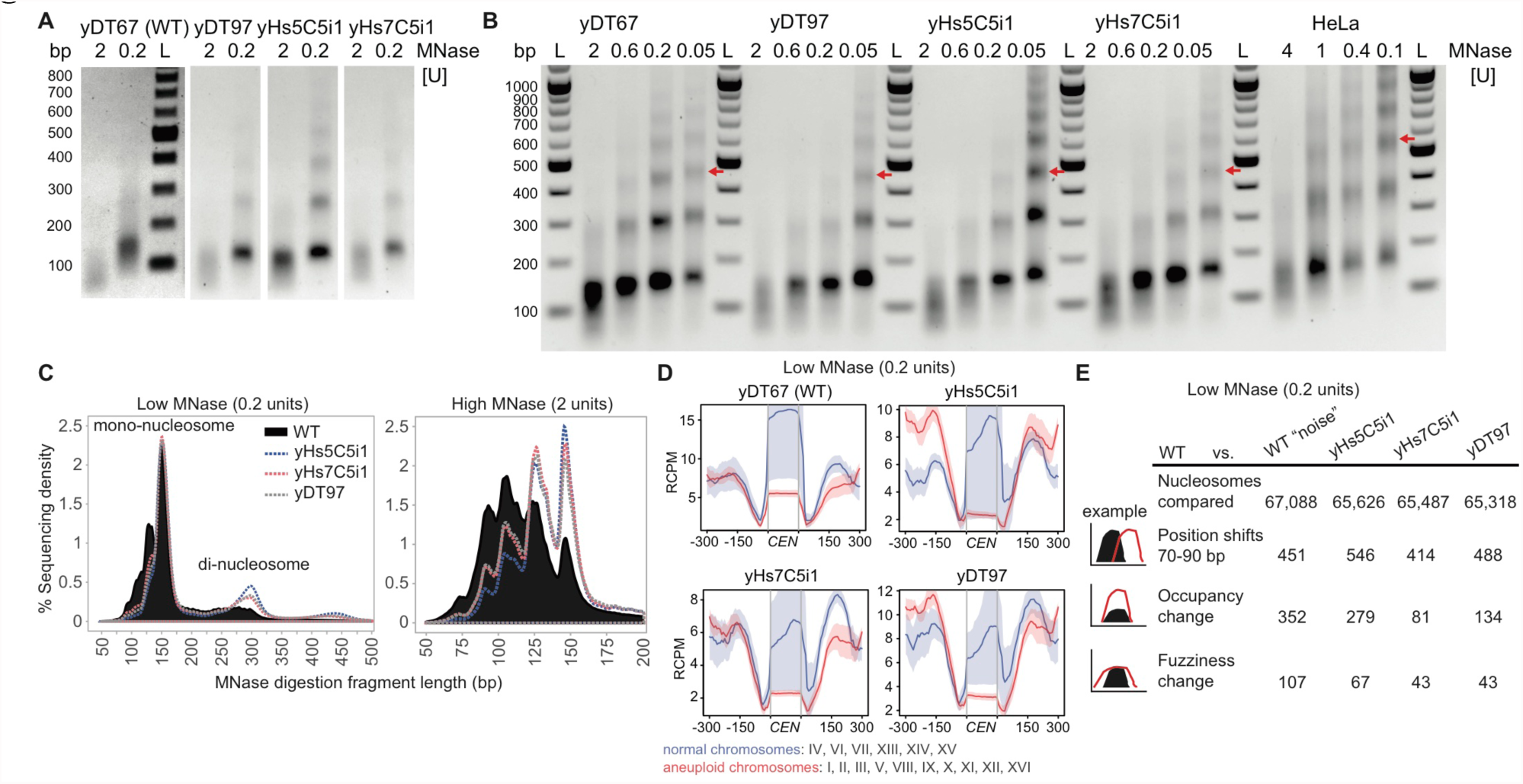
MNase digestions and MNase-seq of humanized yeast, Related to Figure 5. (A) Representative DNA fragments of high (2 units) and low (0.2 units) chromatin MNase digestions used for MNase-sequencing run on a 1% agarose gel. Experiment 1 was performed in biological triplicate and experiment 2 was performed once. All samples from same strain had similar profiles. WT high MNase digests consistently produced a lower DNA yield (3-4 fold), suggesting more accessible DNA, but we chose not to normalize to this because we did not use spike-in controls. “M” refers the DNA marker. (B) Full MNase-titration digestion agarose gel shown in Figure 5A. As WT digests produced less yield, these gels show normalized DNA loading. Red arrows indicate position of the tri-nucleosome, which differs only in the human cell line nucleosome digest. HeLa cells were digested at higher concentrations for a shorter duration and with sonication. “L” refers the DNA marker and “bp” indicates base-pair size. (C) Fragment length histogram from the low and high MNase-seq reads. (D) Low MNase-seq read counts at centromeric regions, plotted for chromosomes that were normal or aneuploid in Figure 2D. RCPM refers to read counts per million mapped reads. (E) Table of Low (0.2 units/ml) MNase-seq nucleosome dynamics between humanized to WT yeast, and WT experiment 1 to WT experiment 2 (“noise”). Occupancy and fuzziness changes use a strict False Discovery Rate cut-off of 0.05 (p < 10^-85^) and additional parameters in *Methods*.

**Figure S7.**
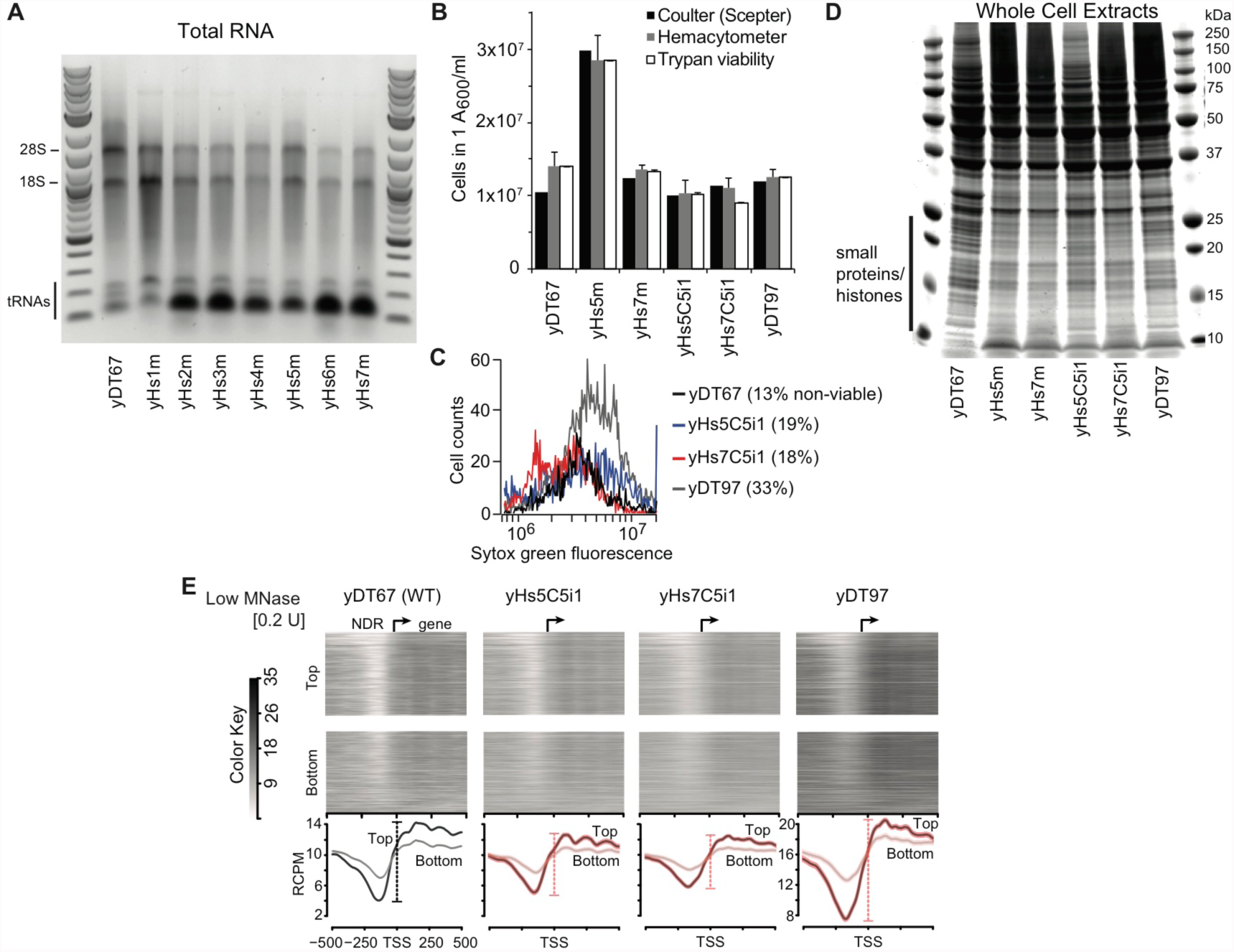
Humanized yeast RNA and protein levels, Related to Figure 6. (A) Total RNA from humanized cells have a similar rRNA to mRNA pattern and ratios as WT cells, although most have elevated tRNA expression. Because tRNA levels are so elevated, mRNA levels are likely lower than indicated in Figure 6A. (B) Reduced RNA content is not due to reduced cell numbers per A_600,_ as yHs cells possess identical or even higher numbers of viable cells (≥10^7^ cells or A_600)._ Cells were measured using both coulter counting (Millipore Scepter) and hemocytometer microscopic counting. Viability was determined by counting number of cells that exclude Trypan blue staining. Bars show standard deviation of 2 replicates. (C) Percentage of non-viable cells (cell viability) determined by Sytox green uptake into dead cells and measured by flow cytometry. (D) Whole-protein extracts of indicated strains run on 12% SDS-bis-Tris acrylamide gel and stained with Coomassie blue. Protein yields were similar on a per cell basis, and each lane has 50 μg total protein loaded. Proteins <25 kDa (e.g., histones) appear reduced in abundance. (E) Heatmaps and average profiles of low concentration MNase-seq reads aligned around the transcription start sites (TSS) ±500 bp of the top and bottom 1500 genes. RCPM refers to read counts per million mapped reads.

**Table S1.**
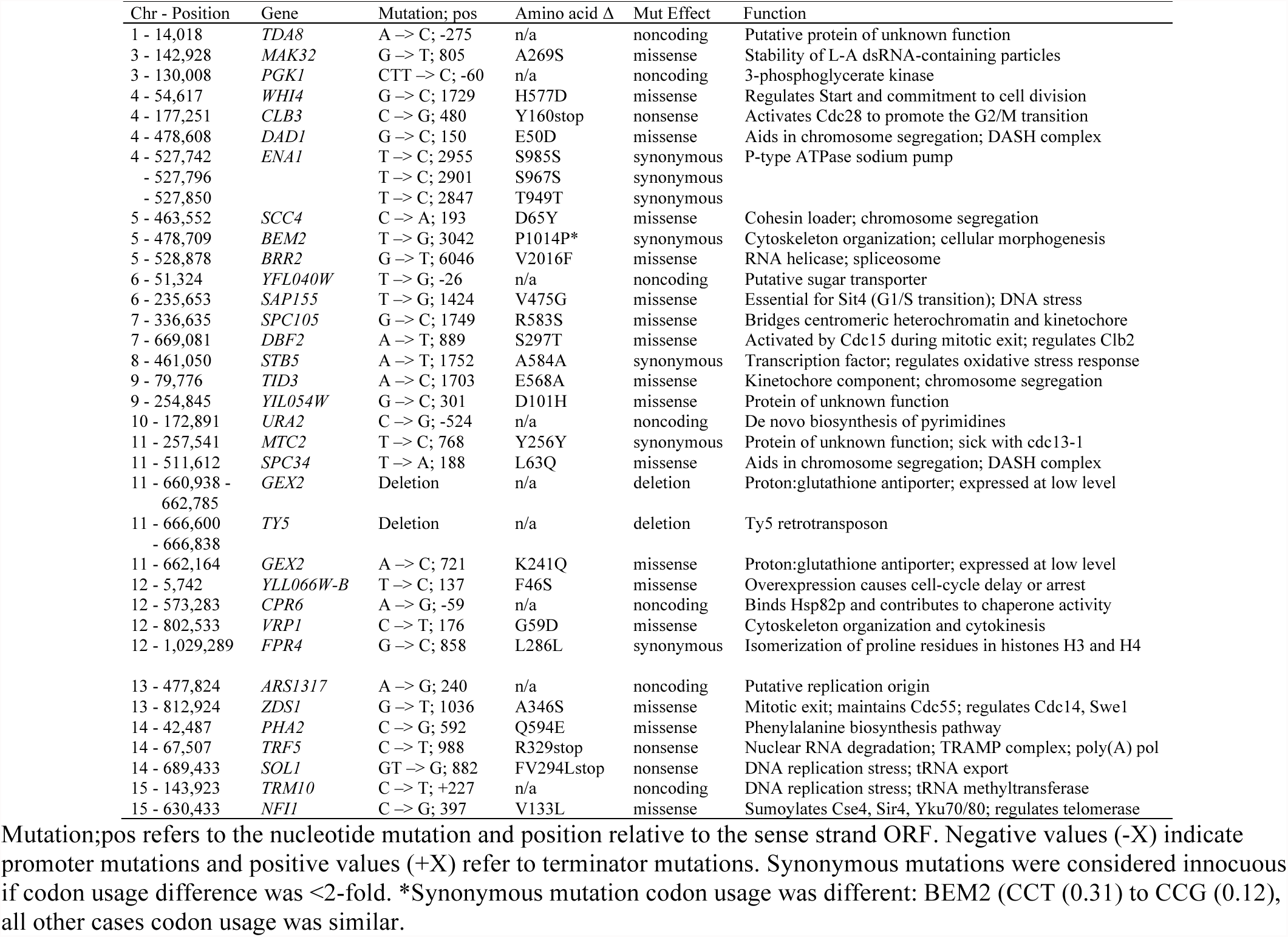
Whole genome sequencing mutations, Related to Figure 2.

**Table S2.**
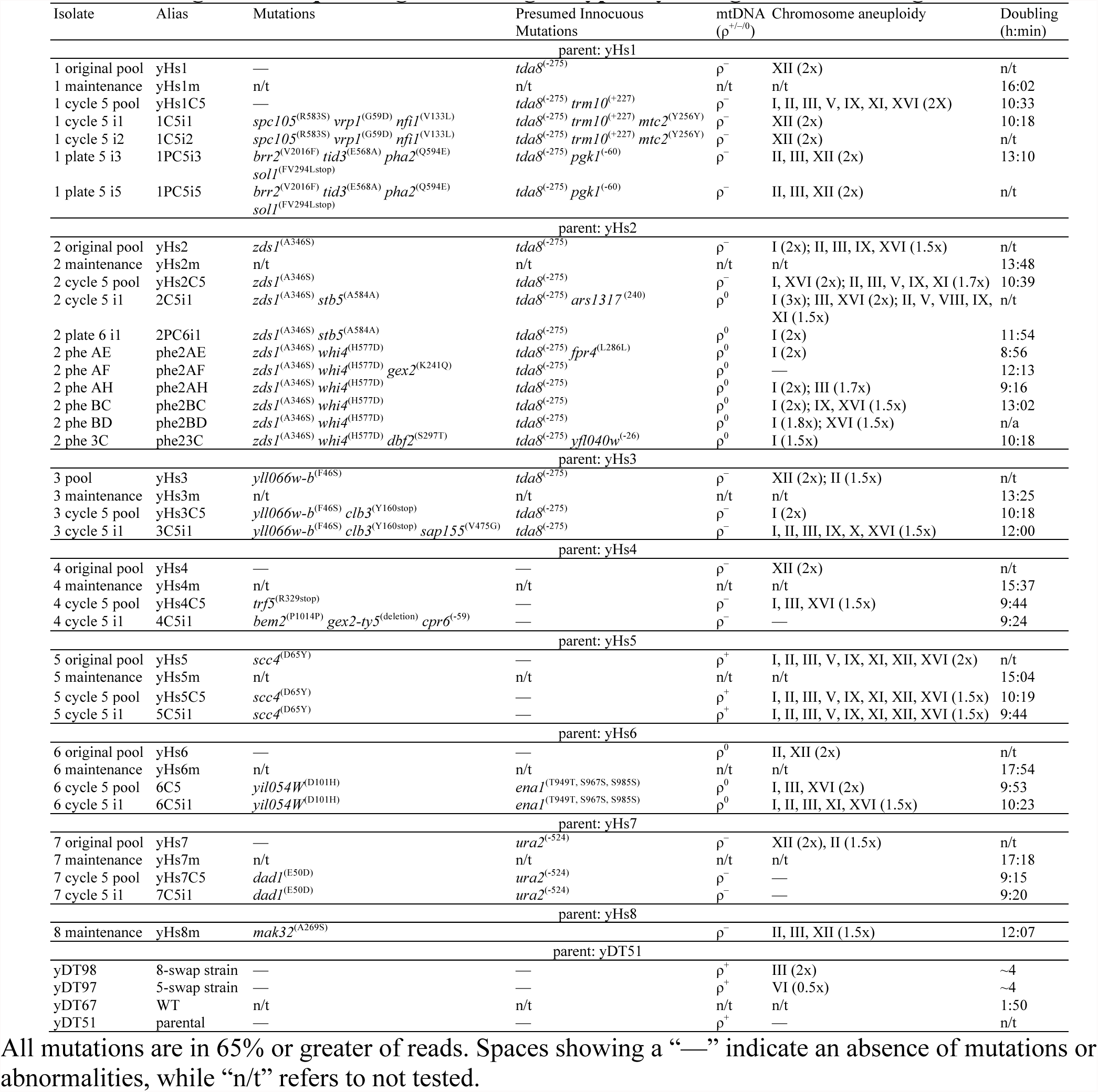
Whole genome sequencing mutation genotypes by lineage, Related to Figure 2.

**Table S3.** Initial humanization trials.

| Strain | Human histones location | Plates (10<sup>7</sup> cells per plate) | Human colonies | Date | Notes |
| --- | --- | --- | --- | --- | --- |
| yDT51 | Low-copy plasmid | 3 | 3 | 04/02/2016 | 15 cm 20 ml plates |
| yDT51 | Low-copy plasmid | 4 | 4 | 05/31/2016 | 15 cm 20 ml plates |
| yDT51 | Low-copy plasmid | 20 | 1 | 06/10/2016 | 15 cm 30 ml plates |
| yDT51 | Genomic integration at *HO*-locus | 10 | 0 | 06/10/2016 | 15 cm 30 ml plates |
| yDT51 | 2-micron High-copy plasmid | 20 | 0 | 06/10/2016 | 15 cm 30 ml plates |
| yDT51 | Low-copy plasmid | 3 | 1 | 11/10/2016 | 15 cm 20 ml plates |

**Table S4.** Evolution cycles from Figure 2A (A_600_ starts at 0.01).

|  | cycle 1 |  | cycle 2 |  | cycle 3 |  | cycle 4 |  | cycle 5 |  |
| --- | --- | --- | --- | --- | --- | --- | --- | --- | --- | --- |
| | $A_{600}$ | days | $A_{600}$ | days | $A_{600}$ | days | $A_{600}$ | days | $A_{600}$ | days |
| yHs1 | 4.0 | 15 | 5.1 | 10 | 6.1 | 8 | 6.6 | 6 | 2.3 | 4 |
| yHs2 | 5.3 | 11 | 6.2 | 6 | 4.2 | 4 | 7.2 | 5 | 5.4 | 4 |
| yHs3 | 6.9 | 11 | 6.2 | 6 | 5.1 | 5 | 6.5 | 5 | 2.5 | 4 |
| yHs4 | 5.2 | 8 | 6.3 | 4 | 6.2 | 4 | 4.9 | 4 | 5.9 | 4 |
| yHs5 | 6.8 | 7 | 4.0 | 4 | 7.1 | 4 | 6.0 | 4 | 6.6 | 4 |
| yHs6 | 5.3 | 10 | 6.3 | 4 | 4.0 | 3 | 6.4 | 3 | 6.3 | 4 |
| yHs7 | 6.0 | 10 | 4.8 | 7 | 5.0 | 4 | 5.3 | 4 | 6.2 | 4 |

**Table S5.**
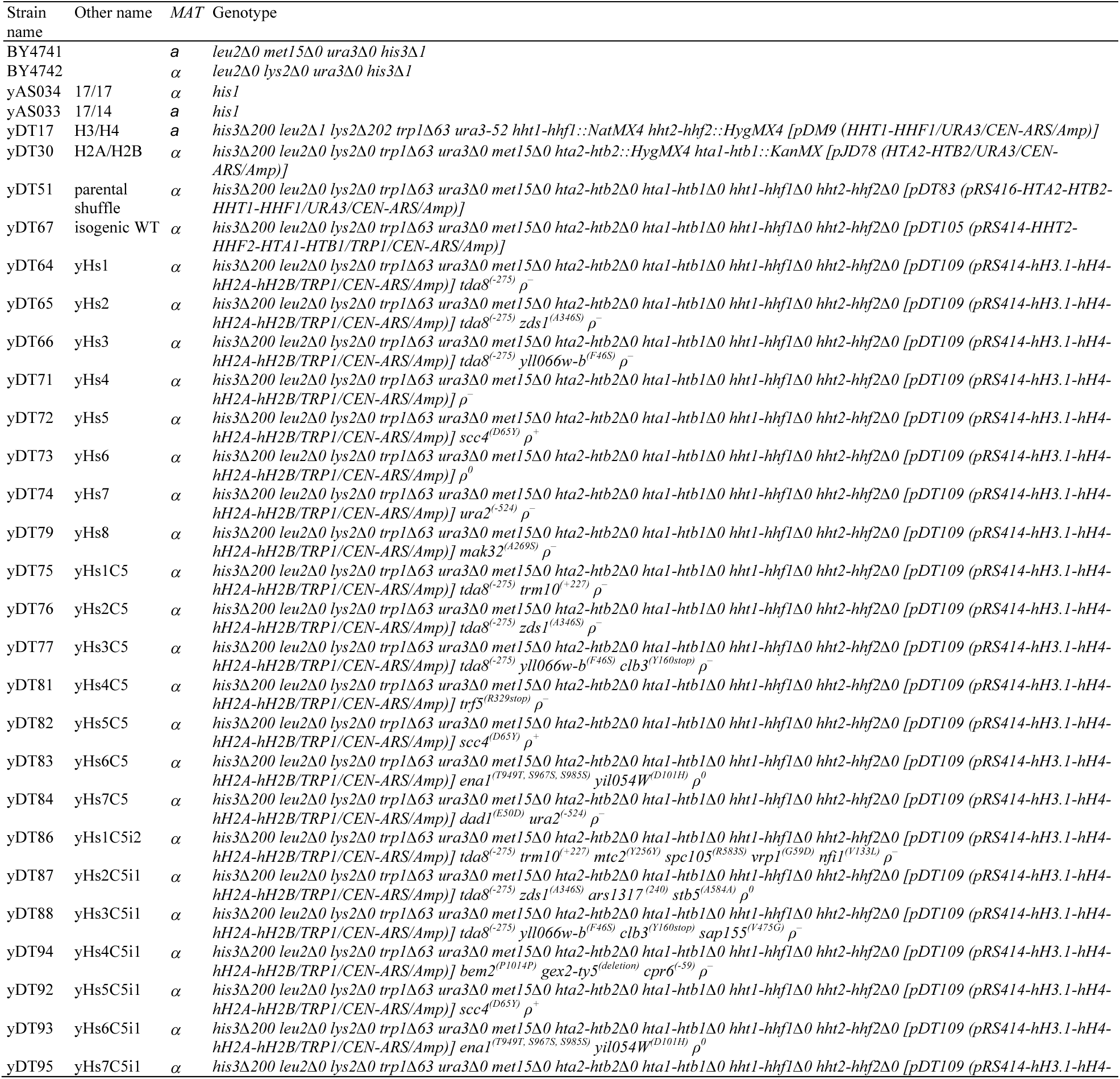
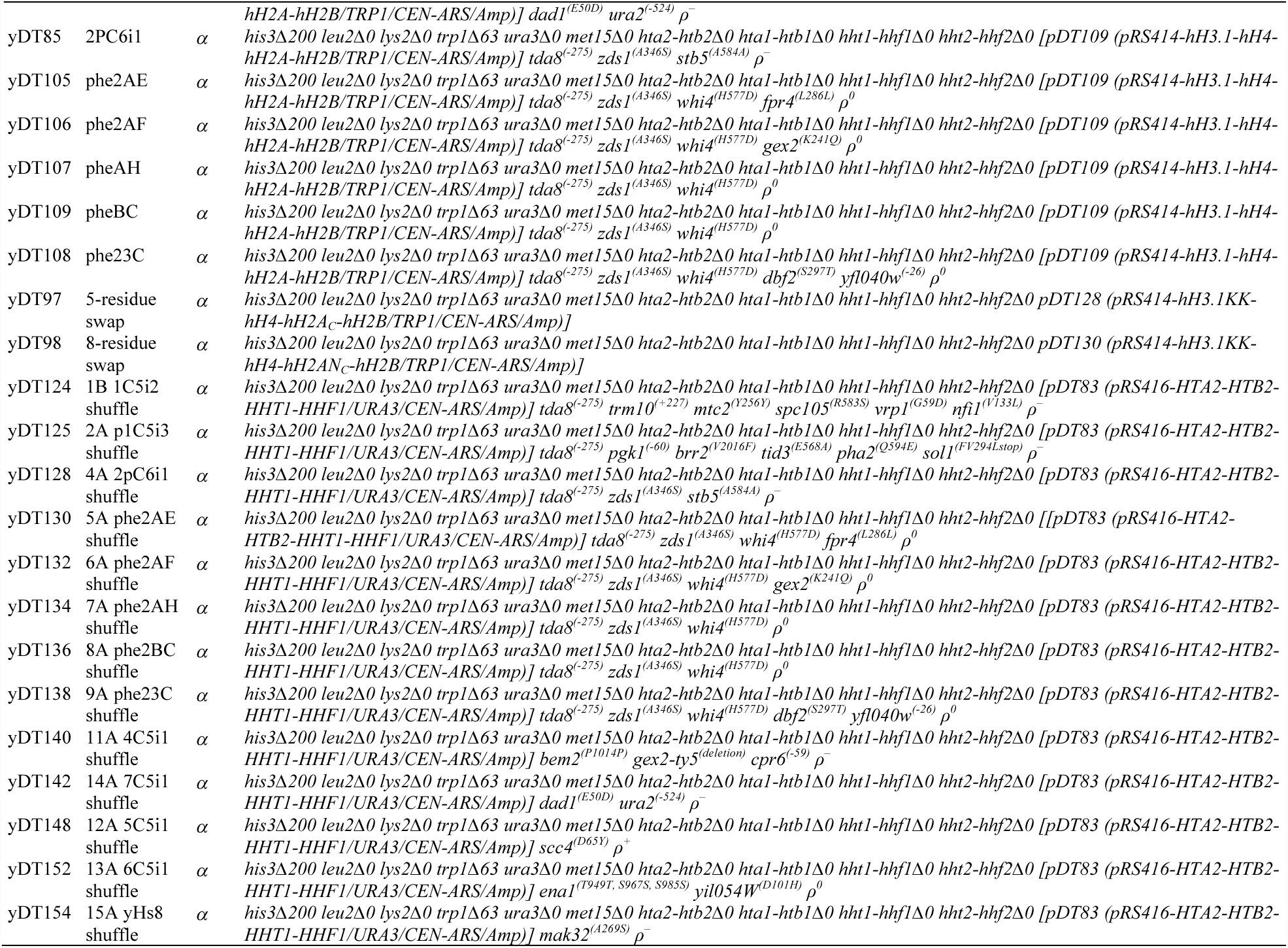
Strains used in this study.

**Table S6.**
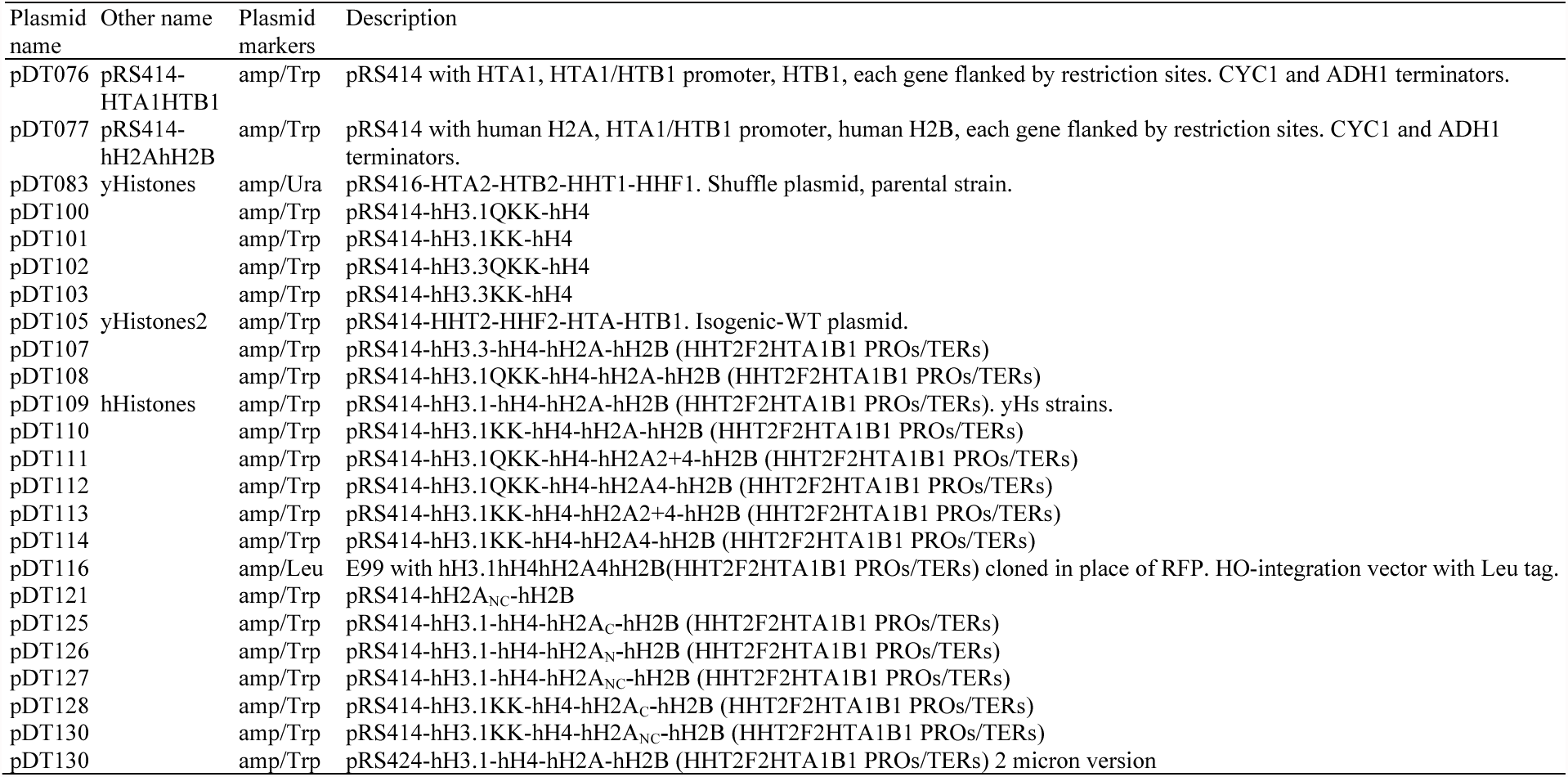
Plasmids.

**Table S7.**
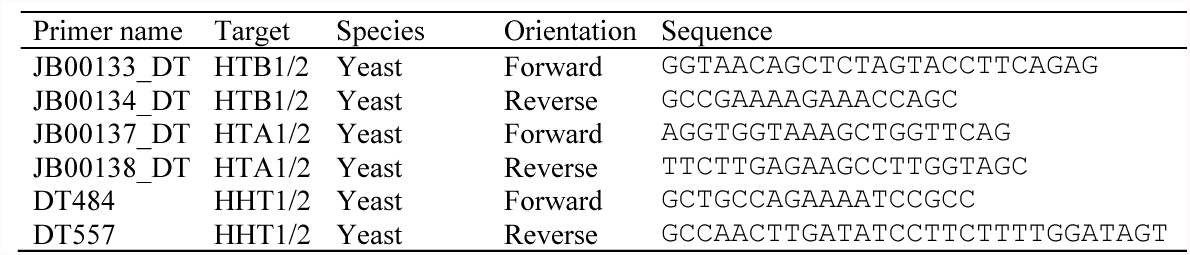
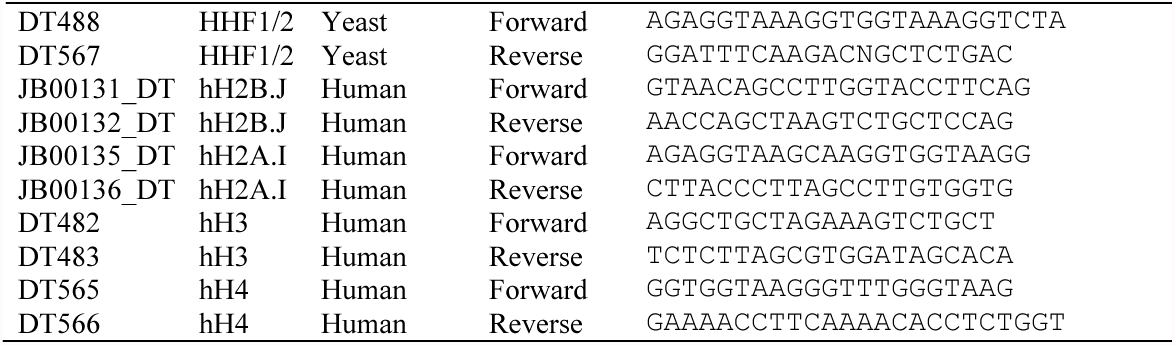
PCRtag primers.

**Table S8.**
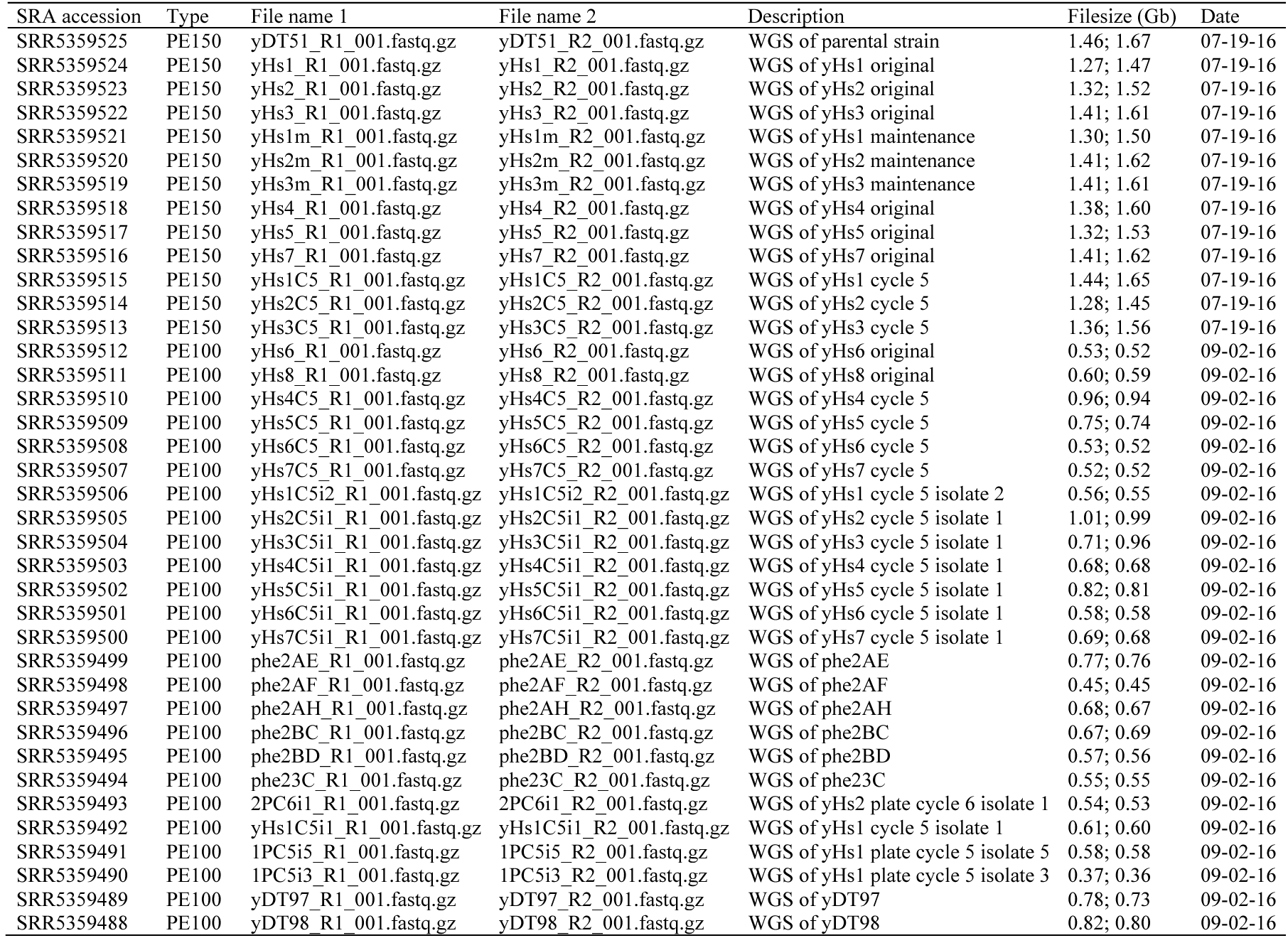
Illumina high-throughput whole-genome sequencing files.

**Table S9.**
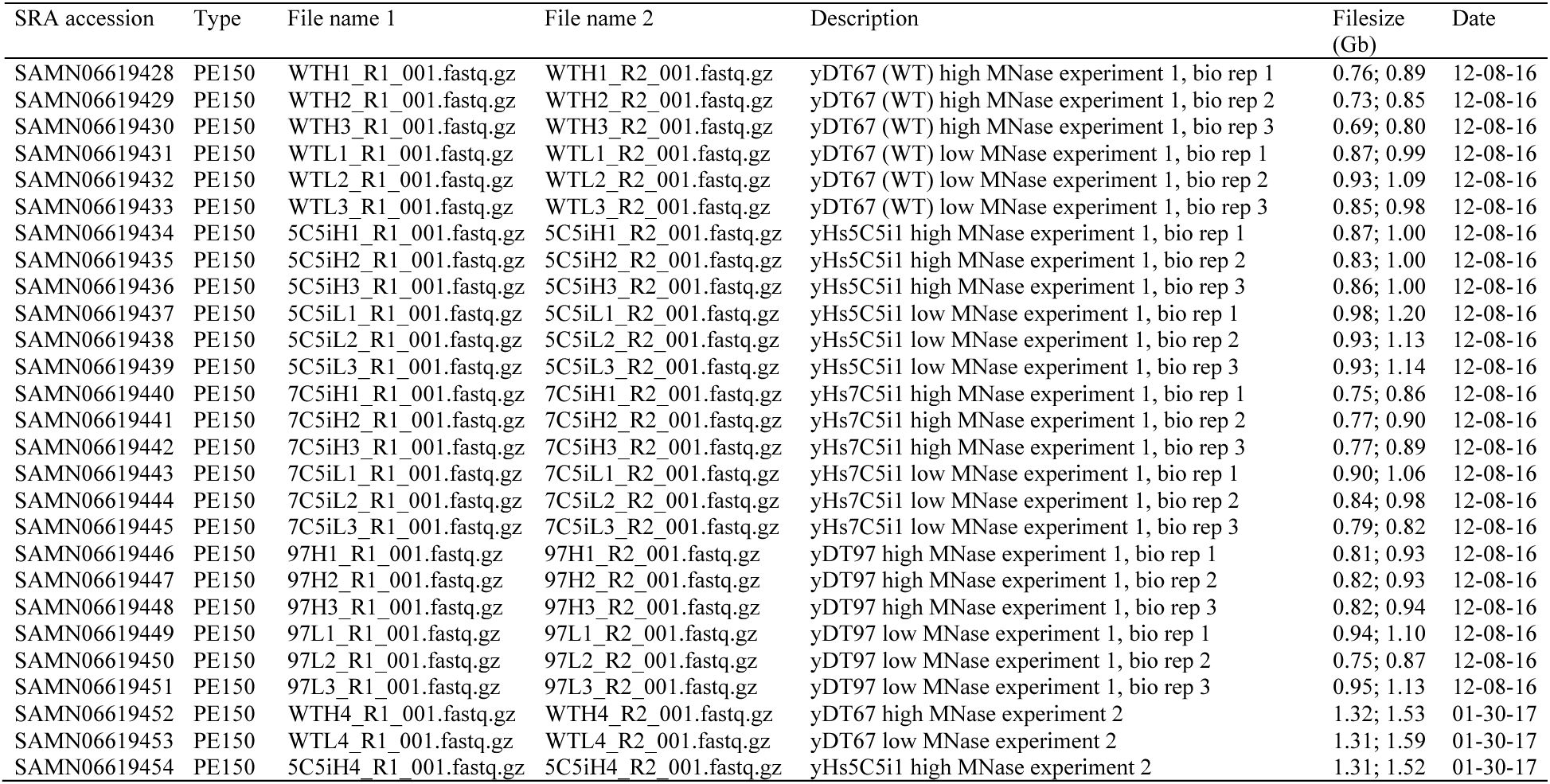
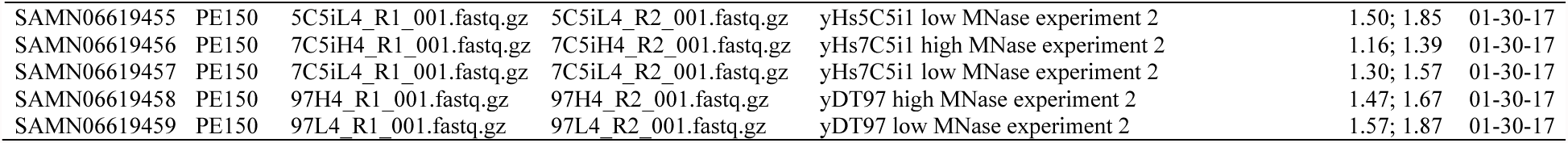
Illumina high-throughput MNase DNA sequencing files.

